# Drought reshapes enhancer-like nascent transcription and gene regulation in *Oryza sativa*

**DOI:** 10.64898/2026.07.14.737839

**Authors:** Fabian Ortner-Krause, Marina Goliasse, John Gitau, Aurore Johary, Farzaneh Rahmdani, Zoé Joly-Lopez

## Abstract

Drought increasingly constrains global rice productivity, yet how water deficit remodels *cis*-regulatory activity in plants remains poorly resolved. Here we used precision run-on sequencing (PRO-seq) to profile nascent transcription in rice leaves under well-watered and drought conditions and mapped transcription-initiation regions with the tool dREG, which detects genome-wide peaks of bidirectional transcription displaying active-enhancer behaviour. PRO-seq captured a robust drought response at genes and revealed extensive remodelling of initiation landscapes. We detected 85,764 consensus dREG sites, of which 17,193 changed significantly under drought and were predominantly intergenic. Because plant intergenic space is rich in transposable elements and silencing-associated transcription, we integrated transposable-element overlap and small-RNA loci with chromatin accessibility and DNA methylation to prioritize 2,428 drought-responsive intergenic sites (841 induced and 1,308 repressed) that are accessible, locally hypomethylated, and bidirectionally transcribed - features consistent with enhancer-like elements. Activity at proximal candidates correlated with elevated nascent transcription of nearby genes, and a subset overlapped gene-connected chromatin loop anchors, supporting candidate enhancer–target relationships. Motif enrichment further supported the involvement of drought-responsive regulatory programs, and hundreds of candidates overlapped rice STARR-seq enhancers. Together, these data define a drought-responsive atlas of candidate enhancer-like nascent transcription in rice and provide prioritized *cis*-regulatory candidates for mechanistic validation and crop improvement.

## Introduction

Drought is now recognized as one of the most severe and fast-growing constraints on global agricultural productivity. Rising temperatures and shifting precipitation patterns have intensified the frequency, duration, and severity of drought events, adding pressure on already strained food systems and threatening food security in vulnerable regions (Bailey-Serres et al., 2019; Park et al., 2025). Among staple crops, rice (*Oryza sativa*) is particularly susceptible due to its high water requirements in irrigated systems and shallow root architecture. Because rice supplies a large share of the daily caloric intake of much of the world’s population, even moderate drought-induced yield penalties carry outsized consequences for global nutrition (Jägermeyr et al., 2021; GNAFC, 2025). Rice faces significant yield losses under water-deficit conditions, making drought resilience a critical breeding target (Gnanamanickam, 2009; Rahman and Zhang, 2023). To maintain resilience under stress, plants activate complex gene-regulatory programs that reconfigure transcriptional landscapes and coordinate adaptive responses (Wilkins et al., 2016; Tripathi and Wilkins, 2021; Hu et al., 2022).

Gene expression in eukaryotes is governed in part by a network of *cis*-regulatory elements (CREs) that orchestrate transcription-factor (TF) binding across various genomic regions, including core promoters and proximal and distal regulatory sequences such as enhancers, silencers, and insulators (Schmitz et al., 2021; Yasmeen et al., 2023; Jores et al., 2025). These elements provide the binding platforms that direct TF recruitment to specific transcription-factor binding sites (TFBS), thereby influencing assembly of the transcriptional machinery and modulating gene activity (Rushton, 2016). Enhancers, which can be located proximal to or distal from their target genes — even thousands of bases to megabases away — are key noncoding DNA sequences that regulate transcription (Wittkopp and Kalay, 2012; Tippens et al., 2018; Yan et al., 2019). They consist of short DNA motifs and control spatiotemporal and cell-type-specific gene expression by interacting with promoter regions to modulate transcriptional output (Buffry et al., 2016; Marand and Schmitz, 2022; Marand et al., 2023). Enhancers have been shown to play key roles in plants’ stress responses (Zhou et al., 2021; Huang et al., 2023).

In animals, active enhancers are most often marked by the production of short, bidirectional, unstable, nonpolyadenylated noncoding transcripts known as enhancer RNAs (eRNAs), which correlate with enhancer activity (Kim et al., 2010; Arnold et al., 2013; Consortium et al., 2014; Core et al., 2014; Arnold et al., 2020). This coupling between transcription and regulatory activity is the conceptual basis for using nascent-transcription assays to identify active enhancers genome-wide: rather than inferring function from static chromatin marks alone, one reads out where engaged RNA polymerase is actively initiating. However, while enhancer transcription is well characterized in mammals, its function in plants remains less defined. A central open question is how faithfully the animal eRNA paradigm transfers to plants, where canonical enhancer marks are less uniform and the boundary between promoters and enhancers may be more fluid (Beernink et al., 2024; McDonald et al., 2024). Emerging evidence nonetheless suggests that bidirectional transcription can mark active enhancers in plants, including crops such as maize and cassava (Ricci et al., 2019; Lozano et al., 2021). Recent work from our group expanded this understanding by identifying thousands of bidirectionally transcribed loci in *O. sativa*, many of which exhibit enhancer-like features: enrichment in accessible chromatin, local hypomethylation, and proximity to highly expressed genes (Joly-Lopez et al., 2020; Goliasse et al., 2025).

In animal systems, stress-responsive enhancer transcription is a well-established mechanism for rapid gene activation (Vihervaara et al., 2017). In plants, however, the chromatin signatures and transcriptional dynamics of these elements, particularly under abiotic stress, remain poorly characterized. A further complication, specific to plant genomes, is that intergenic space is largely transposable-element (TE)-derived and is maintained under silencing by RNA-directed DNA methylation and small RNAs. Bidirectional or pervasive transcription in these regions can therefore arise from silencing-associated processes rather than from *bona fide* regulatory activity (Liu et al., 2018; Chow and Mosher, 2023; Cahn et al., 2024). Any genome-wide search for plant enhancers must therefore explicitly separate regulatory transcription from TE/heterochromatin transcription. This gap is particularly critical for rice, where understanding enhancer activity under stress such as drought could inform breeding strategies for climate resilience.

We selected *O. sativa* as our model due to its agronomic importance, extensive genomic resources, and sensitivity to drought. We used precision run-on sequencing (PRO-seq), a method that uses biotinylated nucleotides to label and capture nascent RNA produced by all engaged polymerases, to map and identify bidirectionally transcribed loci (Core et al., 2014; Mahat et al., 2016). To call bidirectionally transcribed regulatory elements from the run-on signal, we used dREG, a sensitive support-vector-regression method that identifies actively transcribed regulatory elements (Danko et al., 2015; Wang et al., 2019; Wang et al., 2022b). Other approaches have also been used to detect enhancer-like transcripts in plants, including GRO-seq (wheat, Arabidopsis), plaNET-seq (native elongating transcripts in Arabidopsis), and cap-based transcription-start-site (TSS) profiling (csRNA-seq in maize and Arabidopsis) (Zhu et al., 2018; Kindgren et al., 2019; Cahn et al., 2024; McDonald et al., 2024). However, PRO-seq offers single-nucleotide, strand-specific maps of engaged polymerases with minimal dependence on RNA processing or stability, making it particularly well suited for accurate mapping of stress-responsive nascent transcription.

In this study, we define the genome-wide, drought-responsive landscape of enhancer-like nascent transcription in *O. sativa* by calling bidirectionally transcribed loci with PRO-seq/dREG and prioritizing high-confidence candidates through an integrative analysis of chromatin accessibility, DNA-methylation contexts, small RNAs, and TE profiling. We detect sites of bidirectional enhancer-like transcription (eRNAs) that change under drought, prioritize active enhancer-like elements by integrating these omic layers, and test their associations with gene expression and 3D genome contacts. Finally, we examine the sequence architecture of these loci to identify enriched transcription-factor binding motifs. Together, this provides a prioritized, multi-evidence catalogue of candidate drought-responsive regulatory elements in rice.

## Results

### Multi-omics design captures nascent transcription under drought

To test whether we could detect the effect of drought stress on nascent transcription in *O. sativa*, we first profiled *O. sativa* cultivar Azucena plants at the juvenile stage (23 days old) subjected to a moderate drought (50% relative water content) for 10 days alongside well-watered controls, with three biological replicates per condition. PRO-seq generated high-depth, strand-specific maps of engaged RNA polymerases, yielding 10.4–26.2 million trimmed reads for well-watered libraries and 22.9–42.5 million for drought libraries (Table S1). Library quality was consistent across replicates, with enrichment of signal over promoters and gene bodies and balanced representation of forward and reverse strands (Supplementary Fig. S1, S2). Principal-component analysis (PCA) showed a clear separation between well-watered and drought-treated plants along PC1 (97%), indicating that drought was the dominant source of variation; biological replicates within each treatment clustered tightly, while PC2 captured only a minor secondary axis of variation within the drought condition (Supplementary Fig. S3).

We next evaluated whether the rice PRO-seq data exhibited the characteristic features of nascent transcription. Two signatures are diagnostic of engaged, unprocessed transcription: promoter-proximal accumulation of RNA Pol II (pausing) and a high intronic-to-exonic read ratio. We calculated a promoter pause index for each expressed gene and found a broad range of values in both well-watered and drought plants, with most genes showing elevated PRO-seq signal in a promoter-proximal window relative to downstream gene bodies (Supplementary Fig. S4A). A gene-level analysis of pause index (PI) (7,631 genes longer than 5 kb passing PI > 0 filtering) found no individual gene with a statistically significant change between well-watered and drought conditions after multiple-testing correction (per-gene Wilcoxon test, 0 of 4,664 testable genes at padj < 0.05, reflecting the three-replicate design), and the genome-wide PI distributions were closely similar between conditions (median PI 1.8 versus 2.2; Supplementary Fig. S5A), indicating that promoter-proximal pausing is broadly maintained under moderate drought. Nonetheless, a minority of genes showed pronounced changes in pausing, with the strongest responders shown in Supplementary Fig. S5B. In addition, the ratio of intronic to exonic reads was consistently high across libraries, consistent with enrichment for unspliced nascent transcripts rather than mature mRNAs (Supplementary Fig. S4B). Together, the promoter-proximal Pol II accumulation and the intron-rich read distribution confirm that our libraries report engaged, ongoing transcription, providing a robust basis for comparing transcriptional responses between conditions.

To contextualize our results with active-enhancer features and potential regulatory mechanisms, we generated complementary datasets for well-watered and drought conditions. Leaf Assay for Transposase-Accessible Chromatin using sequencing (ATAC-seq) was generated to measure chromatin accessibility, alongside whole-genome bisulfite sequencing (MethylC-seq) for cytosine methylation in all contexts (CG, CHG, CHH) (Tables S6, S7). In addition, we used previously generated small-RNA profiles to mark RNA-directed DNA methylation (RdDM)-linked heterochromatin, and Pore-C maps to chart 3D chromatin contacts (see Methods). These complementary datasets were previously shown to be useful for parsing nascent-transcription changes at genes and dREG sites into a broader regulatory framework (Goliasse et al., 2025).

### A genome-wide catalogue of bidirectionally transcribed elements reveals a distinctive drought-induced dREG profile in *O. sativa*

To assess if and how drought stress reshapes regulatory transcription in *O. sativa*, we called bidirectionally transcribed loci using dREG, which is trained to recognize transcription - initiation regions (TIRs) in nascent-RNA data and to assign a dREG score. This score reflects signal similarity to initiation signatures from a reference set of active enhancers in humans (Chu et al. 2018). Although originally developed and benchmarked in animals, dREG has been successfully applied to plants and recovers enhancer-like bidirectionally transcribed loci (Joly-Lopez et al., 2020; Lozano et al., 2021; Goliasse et al., 2025).

Despite good correlation between PRO-seq replicates (Supplementary Fig. S1, S2), we detected differences in sequencing depth among replicates (Table S1) that could impact the number of dREG peaks called per replicate. We therefore favoured a conservative strategy over the file-merging option usually recommended to maximize dREG discovery in mammals: dREG was run independently for each replicate, and consensus peaks were retained when present in two or more replicates within each condition (Table S2; see Methods). This approach produced 46,101 consensus dREG peaks in well-watered (WW) and 39,770 in drought (Tables S2 and S9). Because 31,898 drought peaks overlap WW peaks, merging these into non-redundant loci across the two conditions yields 85,764 distinct genomic regions, which form the basis for downstream differential testing (see Methods). Of these, 31,898 drought dREGs overlapped WW dREGs, while 7,872 drought-specific and 14,101 WW-specific loci showed no overlap (Fig. 1A). Peak coordinates, scores, and summits for all consensus dREG peaks are provided in Table S9. The final peak sets showed broadly similar distributions of peak length and dREG score in well-watered and drought conditions (Fig. 1B). However, drought-treated plants displayed a modest but detectable shift toward longer peaks and higher dREG scores, suggesting that drought may preferentially engage loci with stronger and somewhat more extended nascent transcription. Meta-profiles of PRO-seq signal around all dREG peaks confirmed a strong, peak-centred enrichment of nascent transcription decaying symmetrically around dREG summits, and higher dREG scores corresponded to progressively stronger bidirectional PRO-seq signal (Fig. 1C).

**Figure 1.**
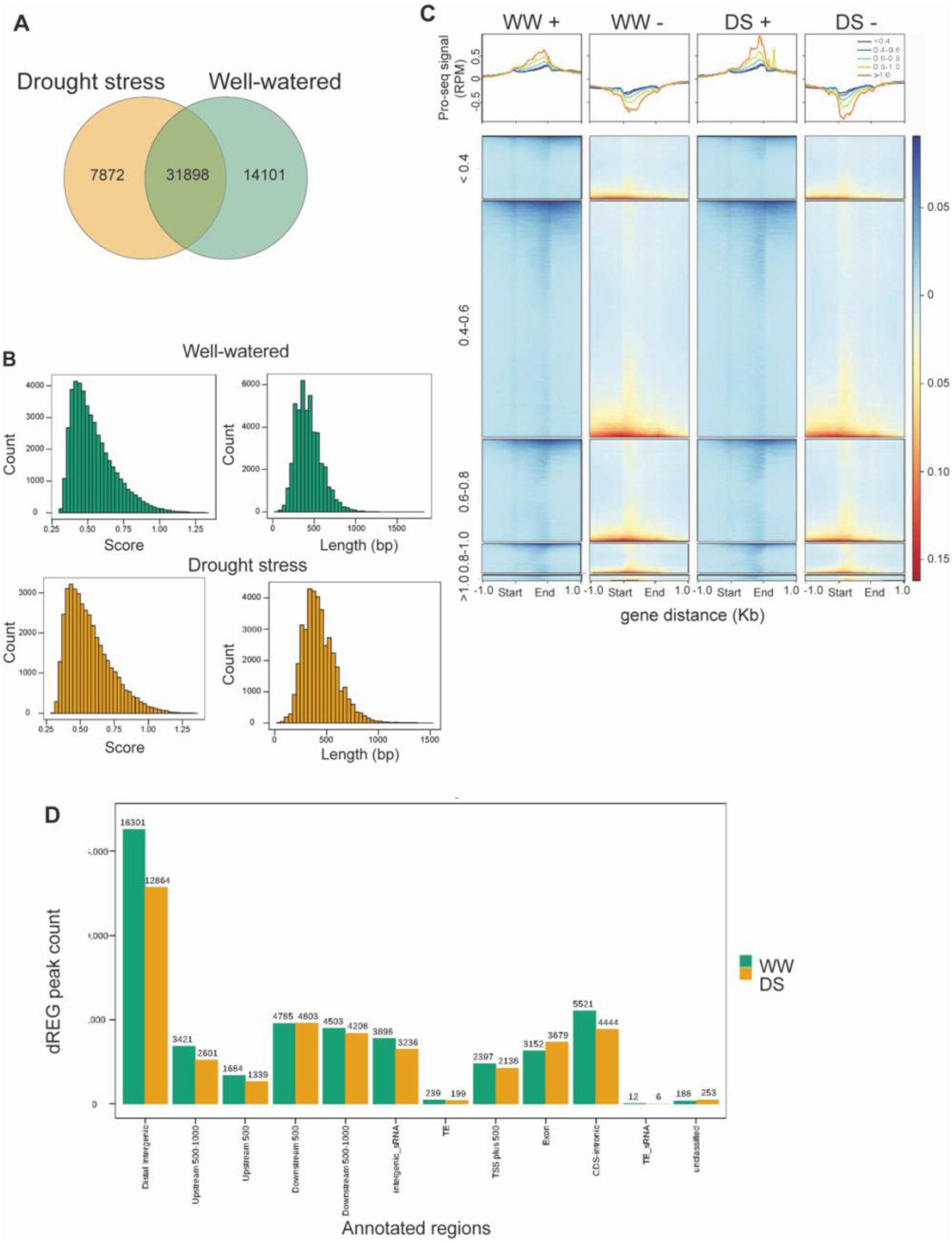
dREG catalogue of bidirectionally transcribed loci. (A) Overlap of consensus dREG peaks between well-watered and drought conditions. (B) Distributions of dREG score and peak length for the final intersected peak sets per condition. (C) Heatmaps and meta-profiles of normalized PRO-seq signal on the plus and minus strands, stratified by dREG-score class. (< 0.4, 0.4-0.6, 0.6-0.8, <0.8-1.0) (D) Genomic distribution of dREG peaks across genic and intergenic features.

Next, we examined the genomic distribution of dREG peaks under both conditions. Genomic annotation placed 21,349 loci in genic regions and 64,081 in intergenic regions (Table S3), indicating that while bidirectional genic initiation is abundant, the majority of divergent initiation occurs proximal and distal to annotated genes (Fig. 1D). Within the genic class, dREG sites were enriched in introns and promoter-proximal windows (TSS ± 500 bp), with comparatively fewer peaks in coding exons, consistent with widespread noncoding transcription initiation within gene bodies. To quantify intronic enrichment more precisely, we intersected condition-level consensus dREG peaks with annotated introns from the Azucena annotation (see Methods). Of the 46,101 well-watered dREGs, 7,474 (16.2%) overlapped annotated introns by at least 1 bp and 3,876 (8.4%) were entirely intronic; similarly, of the 39,770 drought dREGs, 6,544 (16.5%) overlapped introns and 2,916 (7.3%) were fully intronic (Table S3). These proportions indicate that a substantial fraction of genic dREG sites correspond to initiation within introns rather than at annotated exons. PRO-seq profiles centred on intron-overlapping dREG peaks confirmed divergent initiation within intronic sequences (Supplementary Fig. S6). These intronic dREGs may reflect intragenic regulatory elements such as cryptic promoters, alternative transcription start sites, or enhancer-like sequences embedded within gene bodies.

### Drought reshapes gene transcription and reveals that thousands of dREGs are drought-responsive

Because PRO-seq profiles genome-wide nascent transcription, we first confirmed that our data recapitulated the expected genic drought response — that is, changes in active transcription rather than residual steady-state RNA as captured by RNA-seq. We quantified nascent transcription and tested for differentially expressed genes (DEGs) between well-watered and drought using DESeq2 (implemented in the tfTarget framework; (Wang et al., 2022b) on gene-body counts from the PRO-seq data, with padj ≤ 0.05 (Benjamini–Hochberg) and |log2FC| ≥ 1. We identified 2,844 DEGs, including 2,173 upregulated (76.4%) and 671 downregulated (23.6%) in drought (Fig. 2A; Table S4). This DEG count may be lower than reported in some steady-state RNA-seq drought studies in rice; this is expected because gene-body-counted nascent transcription captures actively engaged polymerase rather than accumulated transcripts, and we applied a stringent two-fold cutoff. A volcano plot highlighted numerous strongly up- and downregulated genes across a broad range of effect sizes (Fig. 2B). Gene Ontology (GO) and KEGG pathway analysis (Yu et al., 2012; Wu et al., 2021) indicated enrichment among upregulated genes for defense response, salicylic/jasmonic-acid signaling, leaf senescence, and amino-acid biosynthesis (Supplementary Fig. S7); downregulated genes were enriched for chloroplast organization and cellular responses to metal starvation, although the adjusted p-value was marginally significant owing to the smaller gene set (Supplementary Fig. S8). These responses are consistent with classical moderate-drought experiments in rice (Wehner et al., 2015; Fu et al., 2017; Ali and Baek, 2020; Wang et al., 2020; Awadalla et al., 2024; Munné-Bosch, 2025; Zschiesche et al., 2025) and reinforce that our PRO-seq data capture a robust, physiologically relevant transcriptional response.

**Figure 2.**
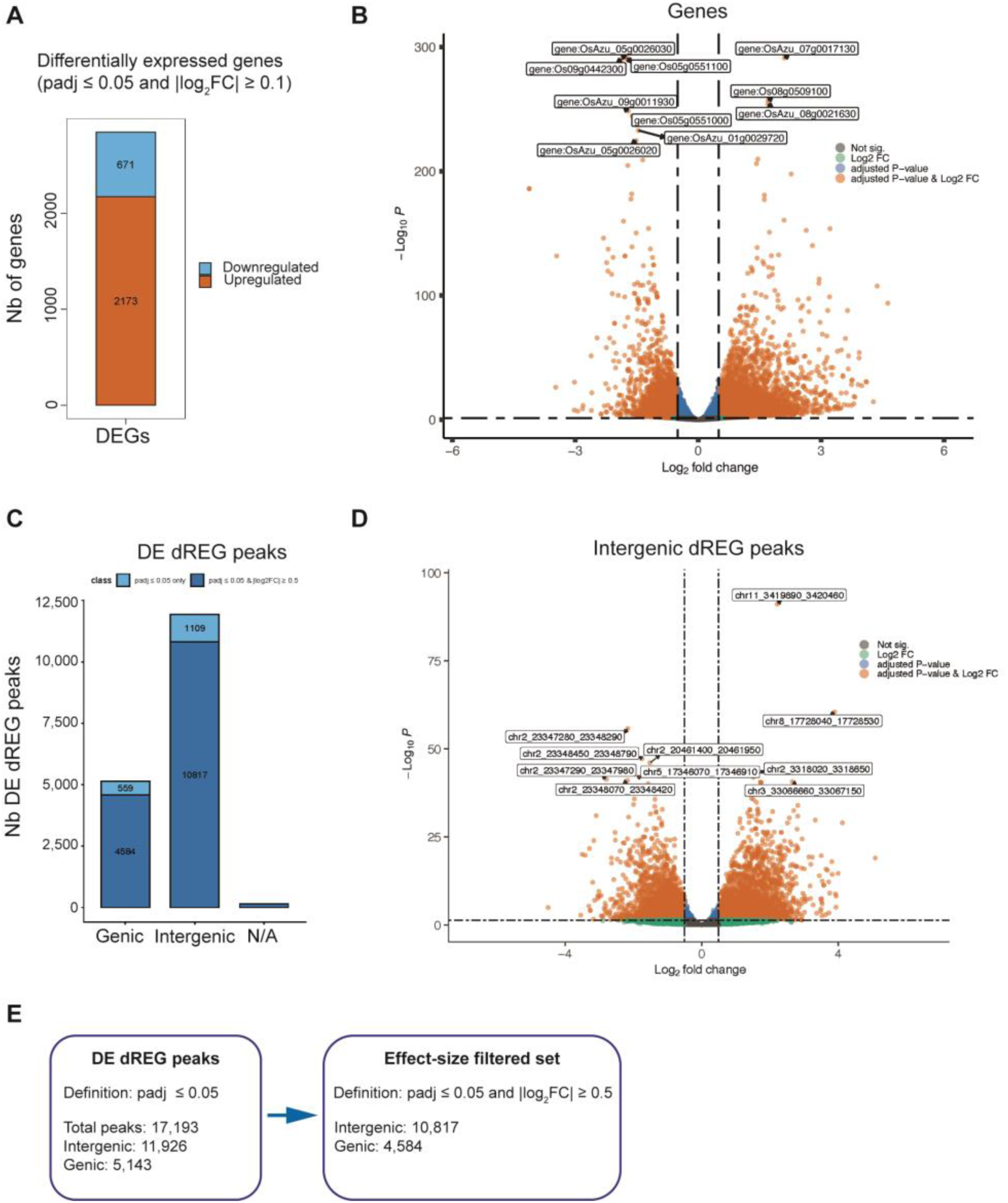
Drought reshapes transcription at genes and dREGs. (A) Numbers of up- and downregulated differentially expressed genes (DEGs (padj ≤ 0.05, |log2FC| ≥ 1). (B) Volcano plot of gene-body nascent transcription. (C) Number of differentially transcribed (DE) dREG peaks (padj ≤ 0.05), split by genic and intergenic annotation. (D) Volcano plot of intergenic dREG peaks. (E) Peak counts retained after the additional |log2FC| ≥ 0.5 effect-size cutoff applied for directionality stratified analyses..

We then asked whether drought similarly reshaped transcription initiation at dREG-defined peaks. Differential calling was performed on the union of condition-level consensus peaks, using tfTarget to quantify PRO-seq signal at dREG peaks and compare conditions. Of 85,764 consensus dREG peaks across both conditions, 17,193 (≈20%) showed significant drought-dependent changes in nascent transcription at padj ≤ 0.05 (Fig. 2C). We then annotated the differentially expressed dREGs using a panel of genomic features — distal intergenic, upstream and downstream regions, gene-internal features, and promoter-proximal regions (TSS ± 100 bp) — and recorded overlap with annotated transposable elements (TEs) and small-RNA (sRNA) loci for subsequent analyses. Peaks overlapping any genic category were classified as genic, whereas those overlapping only distal, upstream, or downstream windows were classified as intergenic. Among these, 11,926 differentially expressed dREGs were intergenic and 5,143 were genic before additional filtering (Fig. 2C). For selected downstream analyses requiring directional stratification, we additionally applied a more stringent effect-size cutoff (|log2FC| ≥ 0.5, padj ≤ 0.05), retaining 10,817 intergenic and 4,584 genic dREGs with robust drought-induced changes in bidirectional transcriptional initiation (Fig. 2E). A dREG-level volcano plot illustrated that these drought-responsive peaks span a wide dynamic range of activation and repression, including many strongly regulated sites that do not coincide with annotated promoters (Fig. 2D). Taken together, drought reshapes nascent transcription not only at genes but also across thousands of potential regulatory elements, the majority of which are intergenic.

### Transposable-element and small-RNA context define a filtered set of enhancer-like intergenic dREGs

Because much of plant intergenic DNA is TE-derived, many intergenic regions are TE-rich, small-RNA-targeted, and heterochromatic. PRO-seq can capture engaged RNA polymerases in these domains as well as at protein-coding genes, promoters, and functional regulatory regions. We therefore asked how TE and small-RNA context influences the behaviour of drought-responsive dREG sites. Using the TE and sRNA annotations recorded for each dREG peak (Fig. 3), we quantified the fraction of each intergenic DE dREG overlapping annotated TEs and whether it overlapped intergenic sRNA loci (Table S5). Intergenic DE dREGs were grouped into TE-overlap bins; most peaks fell into low-to-intermediate TE categories, and median peak length varied only modestly across bins (Fig. 3A). Stratifying peaks by TE-overlap fraction, MethylC-seq profiles showed that DNA methylation increased sharply as TE overlap increased, most markedly in the CHH context, rising steeply beyond ∼20% TE coverage (Fig. 3B). Small-RNA signal followed the same trend and was strongest at highly TE-overlapping peaks (Fig. 3D). Conversely, peaks with little or no TE overlap remained locally hypomethylated and largely free of small-RNA targeting, consistent with a regulatory rather than a silencing origin.

**Figure 3.**
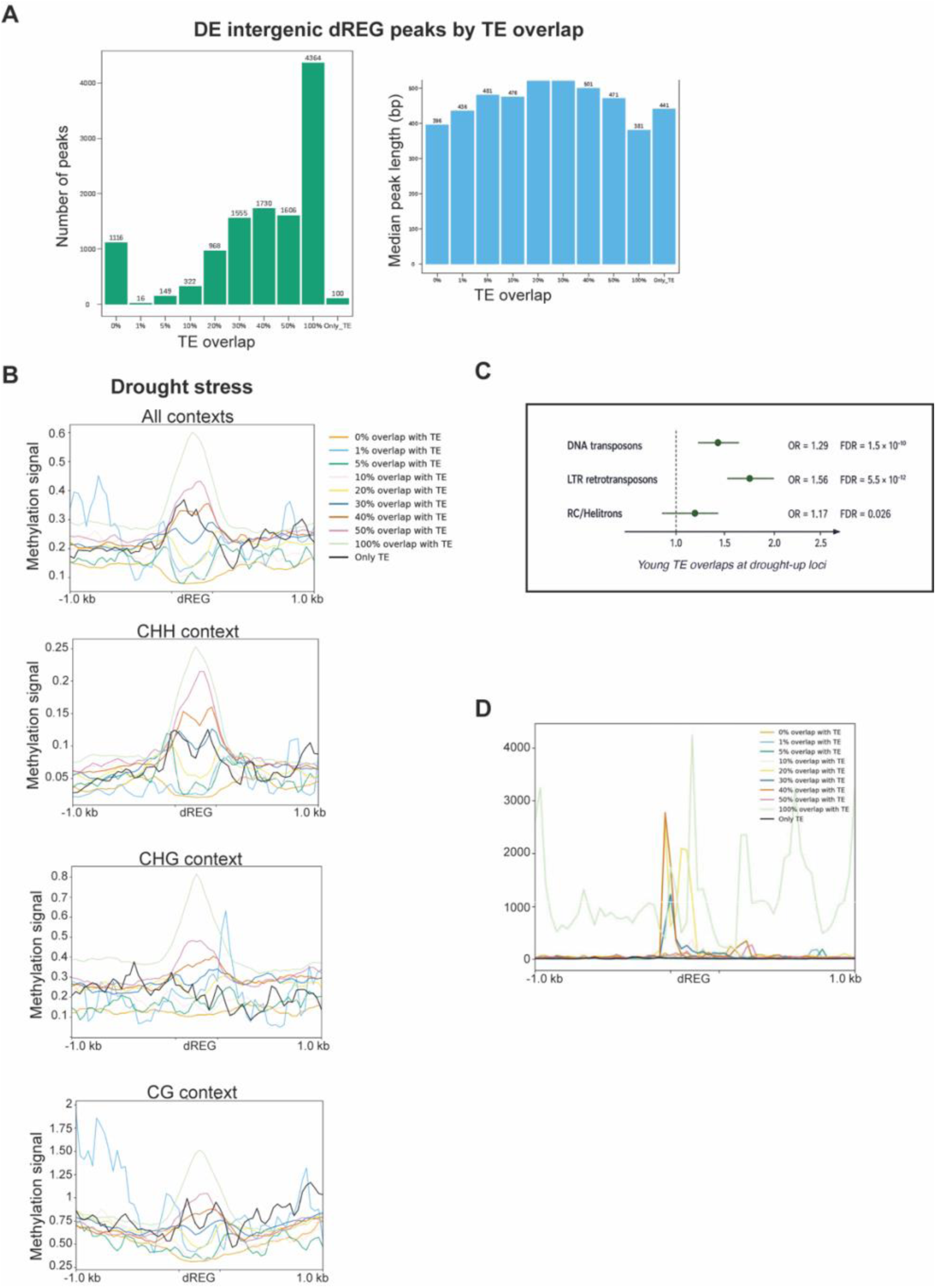
Transposable-element and small-RNA context of intergenic DE dREGs. (A) Distribution of intergenic DE dREG peaks across TE-overlap bins (left) and median peak length per bin (right). (B) MethylC-seq met-profiles within +/− kb od dREG summits, stratified by TE-overlap fraction, shown for all cytosine contexts combined and separately. (C) Odds-ratio enrichment of young transposable-element classes (LTR retrotransposons, DNA transposons, RC/Helitrons) among the 841 drought-upregulated filtered intergenic dREGs, relative to filtered intergenic dREGs passing the same TE/sRNA criteria (two-sided Fisher’s exact test, Benjamini–Hochberg FDR); points show the odds ratio and horizontal lines the 95% confidence interval, with FDR-adjusted significance indicated. (D) Small-RNA signal within ± 1kb of dREG summits, stratified by TE-overlap fraction.

To enrich for regions free of silencing-associated transcription, we considered intergenic dREGs with TE overlap ≤ 20% and no overlap with intergenic sRNA annotations as the most plausible candidates for enhancer-like elements. Starting from 11,926 intergenic dREGs differentially transcribed at padj ≤ 0.05, removing intergenic sRNA-overlapping peaks reduced the set to 10,613 loci, and applying the ≤ 20% TE-overlap cutoff yielded a filtered final set of 2,428 differentially transcribed, unsilenced intergenic dREG sites.

Although the TE/sRNA filter was designed to reduce silencing-associated transcription, it retained a subset of TE-linked drought-responsive candidates. Among young TE-overlapping drought-upregulated loci, we observed significant enrichment for specific TE classes relative to background — most strongly for LTR retrotransposons (odds ratio [OR] = 1.56, FDR = 5.5 × 10⁻¹²) and DNA transposons (OR = 1.29, FDR = 1.5 × 10⁻¹⁰), and more modestly for RC/Helitron elements (OR = 1.17, FDR = 0.026) — suggesting that TE-derived sequences may contribute to a subset of drought-responsive regulatory candidates (Fig. 3C).

### Filtered intergenic dREGs display enhancer hallmarks in chromatin and nascent transcription

While mammalian enhancers are well defined by consistent chromatin signatures and pervasive eRNA transcription, plant enhancers remain far less characterized and may lack these canonical features (Beernink et al., 2024). We therefore asked whether the filtered intergenic dREGs exhibited enhancer-like chromatin signatures. Using within-and-around analyses, we generated meta-profiles and heatmaps of PRO-seq, chromatin accessibility (ATAC-seq), and DNA methylation (MethylC-seq) within ± 1 kb of each filtered DE intergenic dREG, separately for upregulated and downregulated peaks (Fig. 4A). This yielded 841 upregulated and 1,308 downregulated intergenic dREGs (median peak length 410 and 465 bp, respectively; the remaining peaks of the 2,428 padj-filtered set fell below the |log2FC| ≥ 0.5 stratification threshold applied for this directional comparison).

**Figure 4.**
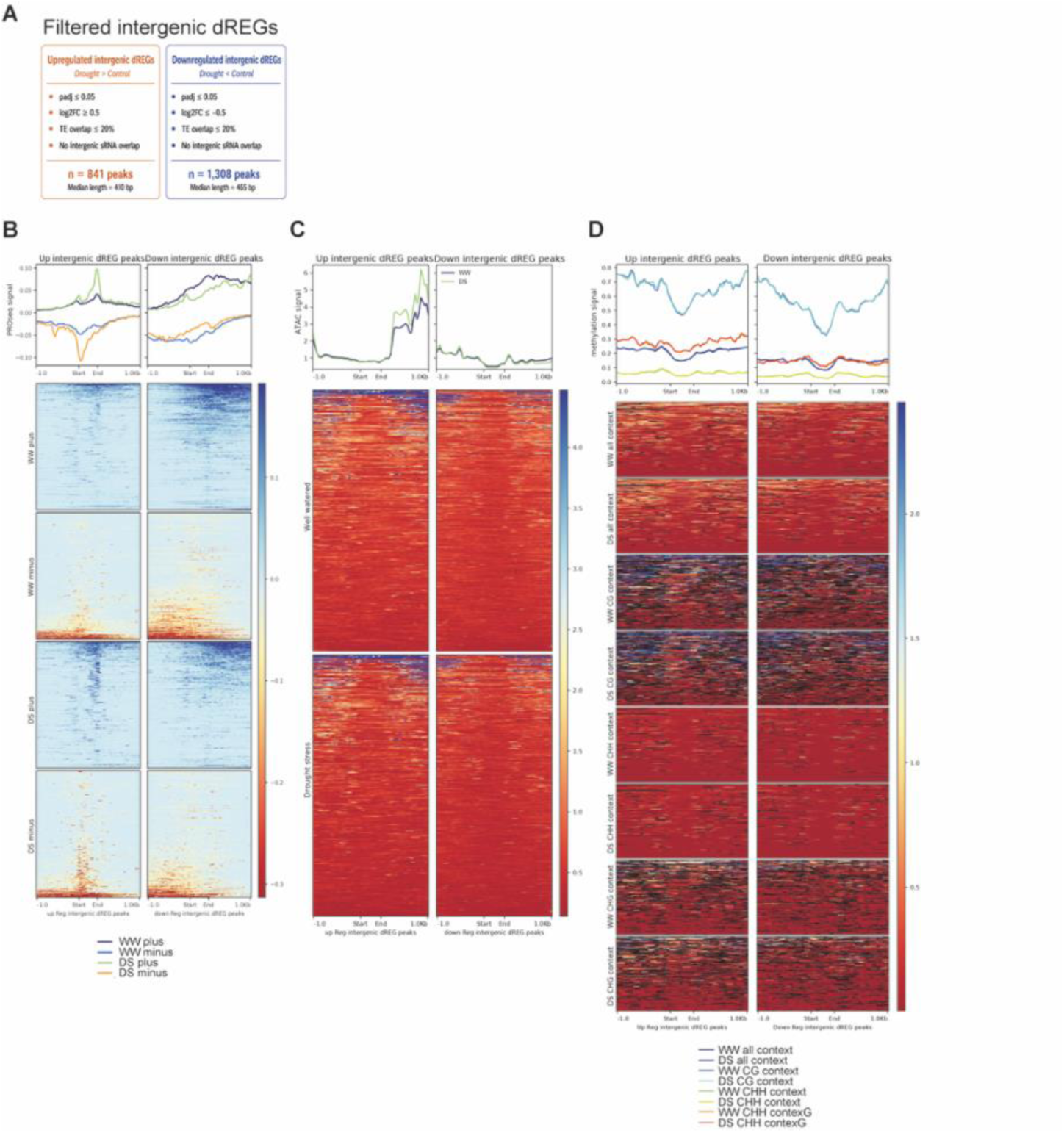
Filtered intergenic dREGs display enhancer hallmarks. (A) Peak-length distribution of 841 upregulated and 1,308 downregulated filtered intergenic dREGs. (B) Within/around PRO-seq meta-profiles showing divergent initiation. (C) ATAC-seq meta-profiles for up-versus downregulated peaks, well-watered versus drought. (D) MethylC-seq meta-profiles showing local hypomethylation across CG and non-CG (CHN) contexts.

PRO-seq signal was strongly enriched over the dREG peaks and was divergent, with plus- and minus-strand reads emanating from opposite sides of the peak centre, confirming the bidirectional initiation captured by dREG (Fig. 4B). Chromatin accessibility was similarly concentrated over the dREG regions, showing a localized ATAC-seq peak with lower accessibility in the flanking sequence (Fig. 4C). Notably, upregulated intergenic dREGs showed higher ATAC-seq signal under drought than under well-watered conditions, consistent with drought-induced chromatin opening at these sites, whereas downregulated intergenic dREGs showed lower overall accessibility with minimal difference between conditions (Fig. 4C). DNA methylation showed a corresponding local decrease at the dREG peak relative to flanking regions across CG and non-CG (CHN) contexts, consistent with local hypomethylation (Fig. 4D). In both peak classes, the dREG centre remained more accessible and less methylated than the surrounding chromatin regardless of the direction of transcriptional change. Taken together, these profiles indicate that the filtered intergenic dREGs collectively exhibit the chromatin and transcription features expected of plant enhancer-like elements in rice.

### Proximal and genic dREGs coincide with elevated gene expression

To explore the enhancer potential of the filtered dREGs, we connected dREG activity with gene output by examining the relationship between differentially expressed dREGs and nearby gene expression. From the filtered intergenic DE dREG set (2,428 peaks), we defined “proximal” as peaks within 1 kb upstream of a transcription start site (TSS) or 1 kb downstream of a transcription end site (TES), using strand-aware annotations from the Azucena gene models. We identified 1,904 proximal intergenic peaks. Intersection with annotated gene coordinates showed that 861 genes harboured at least one proximal DE intergenic dREG, whereas the remaining genes had none in our dataset. The distribution of proximal peak counts per gene was skewed toward single associations, with most genes carrying one or two proximal dREGs (Fig. 5A).

**Figure 5.**
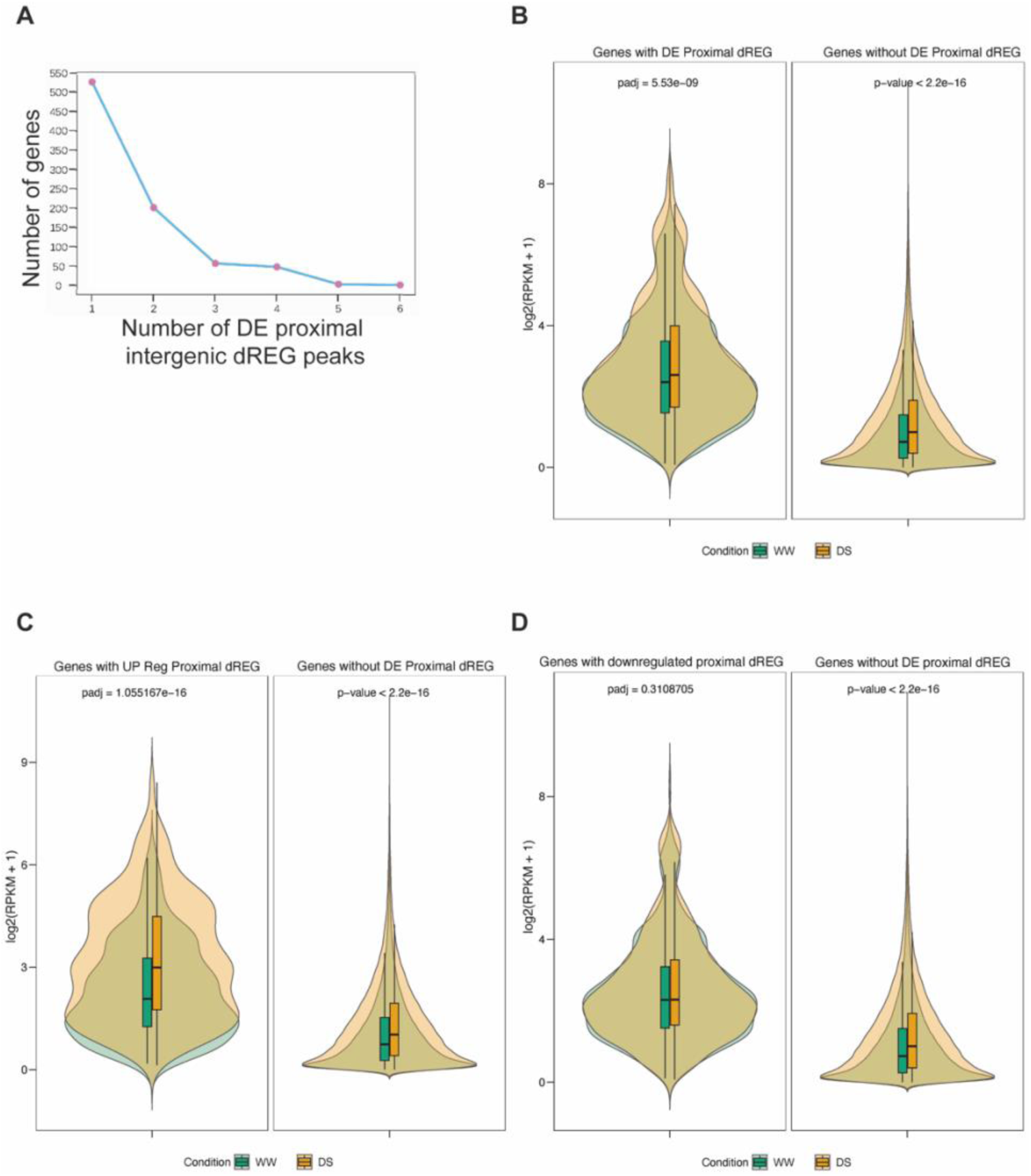
Proximal dREGs coincide with elevated gene expression. (A) Proximal DE dREG peaks per gene. (B) Gene-body expression for genes with versus without a proximal DE dREG. (C, D) Expression stratified by direction of dREG regulation.

We then compared PRO-seq gene-body expression between genes with and without proximal DE dREGs. Using normalized log2 values for each gene and condition, genes harbouring at least one proximal DE intergenic dREG showed higher expression than genes lacking one in both well-watered and drought (Fig. 5B). When stratified by the direction of dREG regulation, genes associated with upregulated proximal intergenic dREGs showed significantly higher expression under drought than under well-watered conditions (padj = 1.06 × 10⁻¹⁶) and were more highly expressed than genes without proximal DE dREGs in both conditions (Fig. 5C,D). In contrast, genes linked to downregulated proximal dREGs did not differ significantly between conditions (padj = 0.31) but remained more highly expressed overall than genes lacking proximal DE dREGs (Fig. 5D). This pattern indicates that the presence of a proximal DE intergenic dREG marks a more highly expressed gene regardless of the direction of the dREG change, while only upregulated dREGs are associated with a condition-dependent increase in nearby gene expression under drought. Similar trends were observed for genic dREGs: genes with differentially expressed genic dREGs showed higher transcription than genes without, and genes with upregulated genic dREGs showed a significant drought-induced increase (padj = 7.80 × 10⁻¹⁹⁴), whereas genes with downregulated genic dREGs did not differ significantly between conditions (padj = 0.35) (Supplementary Fig. S9). These analyses support a model in which drought-responsive dREG activity, both proximal intergenic and genic, is associated with elevated gene transcription, and in which the drought-induced gain in dREG transcription at upregulated sites may contribute to the coordinated activation of nearby genes.

### 3D contacts nominate candidate target genes for intergenic dREGs

Because many intergenic regulatory elements contact their target genes through higher-order chromatin structure, we examined whether the filtered dREGs were embedded in 3D chromatin loop contacts. Using previously published Pore-C maps generated for Azucena leaves (Goliasse et al., 2025); generated under well-watered conditions), we overlaid filtered intergenic dREGs onto chromatin loop anchors and asked whether they formed loops with gene-containing anchors. Filtered intergenic dREGs fell within loop anchors contacting gene bodies or promoters more frequently than expected for length- and distance-matched control regions (3.19-fold enrichment; permutation test, p < 0.001, 1,000 iterations), indicating enrichment of dREGs at gene-connected loop anchors. Restricting to filtered intergenic dREGs that were differentially transcribed under drought, 73 dREG peaks fell within loop anchors contacting gene-bearing anchors, forming 93 dREG–gene loop contacts and linking to 86 candidate target genes. These partitioned strongly by the direction of dREG regulation: 60 upregulated dREG peaks contacted 70 genes, whereas 13 downregulated peaks contacted 16 genes (Fig. 6A). All contacts were intra-chromosomal, and many dREGs contacted genes well beyond their immediate neighbourhood, consistent with long-range rather than strictly promoter-proximal regulatory interactions. This positional association was matched by a directional effect on target-gene output. For each gene–intergenic loop, we compared the drought-induced change in nascent transcription of the gene at the genic anchor according to the dREG present at the paired non-genic anchor. Genes looped to an upregulated dREG showed a significant increase in transcription under drought, whereas genes looped to a downregulated dREG showed a decrease; both differed significantly from genes whose paired anchor carried no differentially transcribed dREG, and the two directional classes differed strongly from one another (Fig. 6B). The dREG at the distal anchor therefore predicts not only whether the looped gene responds to drought, but in which direction, which is an association that is difficult to explain by proximity alone, since all contacts were intra-chromosomal and many spanned distances well beyond the immediate gene neighbourhood.

**Figure 6.**
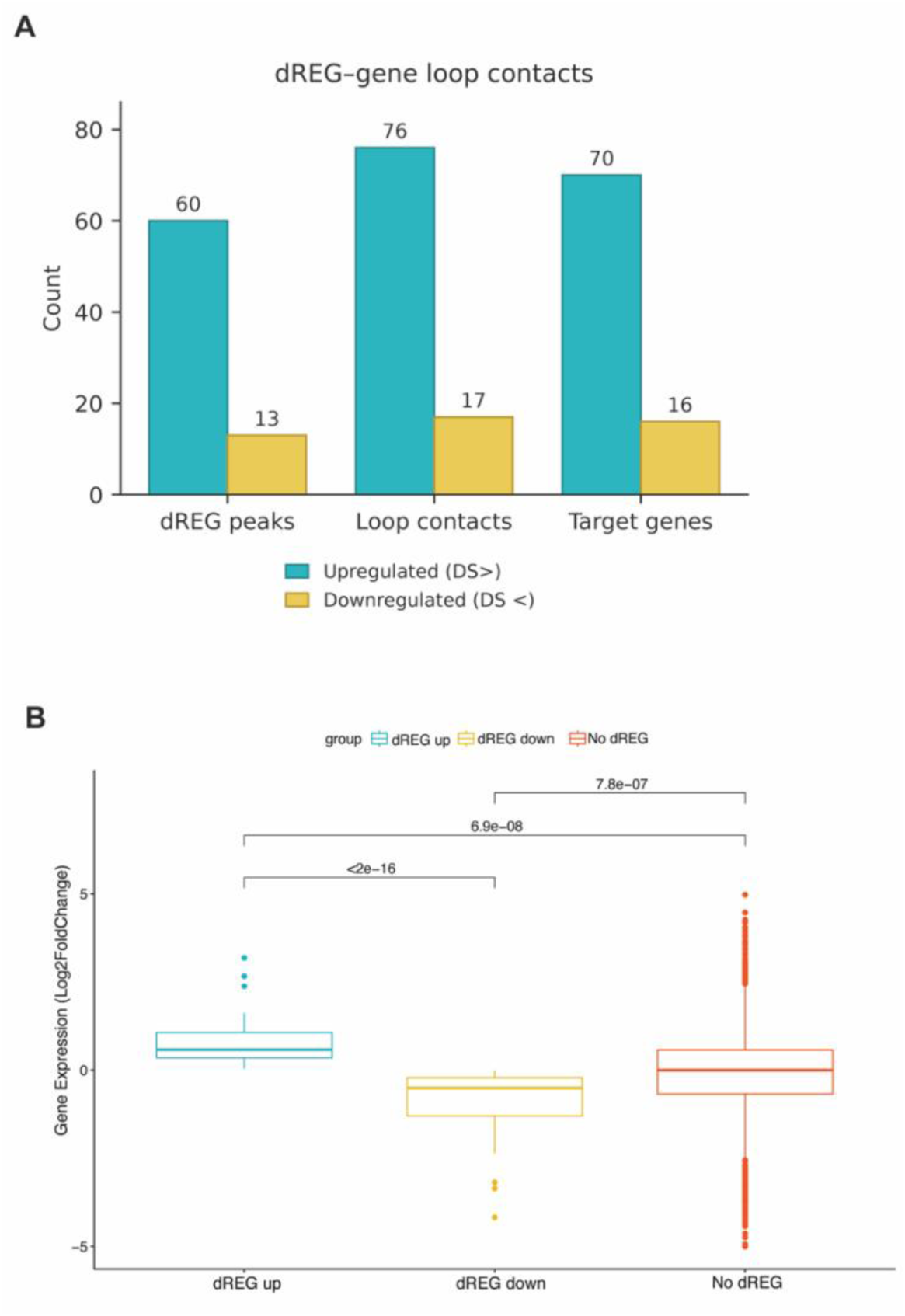
Drought-responsive dREGs at gene-connected 3D contacts are coupled to target-gene transcription. (A) Numbers of drought-DE filtered intergenic dREG peaks located in the non-genic anchor of a gene–intergenic Pore-C loop, together with the resulting loop contacts and candidate target genes, split by direction of dREG regulation. (B) Change in nascent transcription of the gene at the genic anchor (log_2_ fold-change, drought versus well-watered), grouped by the dREG present at the paired non-genic anchor: upregulated (dREG up), downregulated (dREG down), or no differentially transcribed dREG (No dREG). Boxes show the median and interquartile range; whiskers extend to 1.5 × IQR. Pairwise comparisons by two-sided Wilcoxon rank-sum test.

The pronounced predominance of upregulated dREG–gene loops mirrors the activation bias seen at proximal dREGs (Fig. 5) and suggests that at least some intergenic dREG–gene pairs participate in drought-responsive regulatory interactions. Because the Pore-C maps derive from well-watered tissue, these loops indicate which drought-responsive dREGs are positioned within gene-connected 3D contacts rather than drought-induced looping *per se*; future Pore-C or other 3D approaches under drought will help refine these dREG–gene contacts.

### Motif enrichment implicates drought-responsive transcription factors

Having defined a high-confidence set of drought-induced enhancer-like loci, we investigated whether their sequences carry enriched transcription-factor binding motifs that could point to upstream regulatory programs. We performed *de novo* and known-motif enrichment analyses with HOMER on filtered upregulated intergenic dREG peaks (log2FC ≥ 0.5; padj ≤ 0.05; n = 841; 6–8 bp motifs). Relative to a genome-matched background, HOMER recovered multiple significantly enriched motifs (Fig. 7A,B; Supplementary Fig. S10; Table S8), including known matches to TF families broadly associated with stress- and hormone-responsive regulation in plants (e.g., NAC/NAM, bHLH, SBP/SPL, C2H2, and DOF/WRKY-like signatures). These enrichments support the idea that drought-induced intergenic dREG sites contain binding potential for stress-responsive TF programs, while noting that motif enrichment indicates sequence compatibility rather than direct TF occupancy.

**Figure 7.**
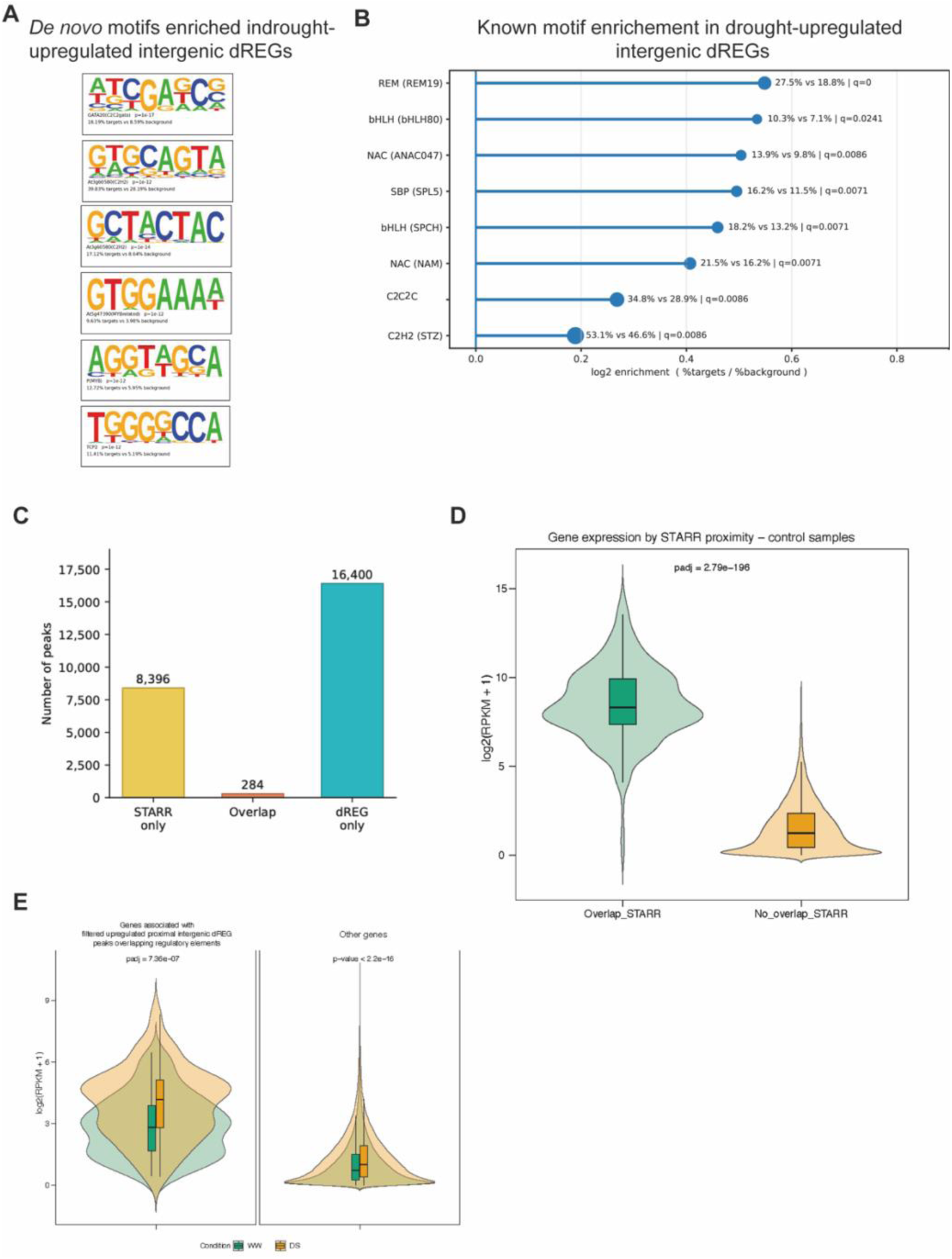
Sequence determinants and orthogonal validation. (A) De novo motifs enriched in the 841 drought-upregulated filtered intergenic dREGs (HOMER). (B) Known-motif enrichment for the same peaks. (C) Overlap between rice STARR-seq enhancers (8,680, lifted to Azucena) and the filtered dREG catalogue (16,828): 8,396 STARR-only, 284 overlapping, and 16,400 dREG-only peaks; the 284 overlapping enhancers correspond to 428 dREG regions. (D) Nascent transcription of genes proximal to a STARR-overlapping dREG (n = 312) versus genes without one. (E) Expression across conditions for genes proximal to filtered upregulated intergenic dREGs overlapping an independent regulatory-element compendium (45 genes; well-watered versus drought).

When we stratified drought-upregulated intergenic dREG peaks by proximity to genes, the overall motif landscape was broadly similar between proximal and distal classes, with several motifs shared by both groups (Supplementary Fig. S11). However, the strength and statistical support of known-motif enrichments differed: the proximal set retained multiple motifs significant after multiple-testing correction, whereas the distal set yielded fewer significant matches. Overall, these patterns are consistent with activated intergenic dREG sites being sequence-compatible with binding by stress- and hormone-responsive TF families, while again noting that motif enrichment alone does not demonstrate TF occupancy.

Finally, we asked whether our filtered dREG peaks coincide with functionally defined candidate enhancers from reporter-based assays such as STARR-seq, which provide an orthogonal readout of autonomous enhancer activity. We used a published rice protoplast STARR-seq enhancer set of 9,642 candidate elements (Sun et al., 2019). Because these coordinates are defined on the Nipponbare reference whereas our dREGs are in Azucena, we lifted the STARR enhancers onto the Azucena assembly and retained 8,680 enhancers for comparison. Intersecting these with our TE/sRNA-filtered dREG catalogue (all genic plus filtered intergenic peaks, n = 16,828) within a ± 200 bp window around each dREG summit (≥ 1 bp overlap), 284 STARR-seq enhancers overlapped dREG peaks — corresponding to 428 dREG regions, reflecting one-to-many STARR–dREG relationships — whereas 8,396 STARR enhancers and 16,400 dREGs were non-overlapping (Fig. 7C). PRO-seq signal at these 284 STARR-overlapping dREGs was strongly bidirectional, and the degree of bidirectionality differed between conditions (paired Wilcoxon test, p = 4.6 × 10⁻¹³), supporting active divergent transcription at these reporter-defined enhancers. Of the 284 overlapping regions, 203 were gene-proximal and were associated with 312 genes; these genes were markedly more highly expressed than genes lacking a proximal STARR-overlapping dREG (Fig. 7D), an effect evident from the shift in expression distributions, although the very large background set inflates the nominal significance.

As an additional and independent line of evidence, we intersected our filtered upregulated proximal intergenic dREGs with a separate, larger published compendium of candidate rice regulatory elements (47,541 sites; (Zhu et al., 2024), distinct from the STARR-seq enhancers analysed above. This identified 176 overlapping peaks associated with 45 genes. Expression of these 45 genes differed significantly with condition (Wilcoxon padj = 7.36 × 10 ⁻⁷), and they were more highly expressed than the remaining genes in both conditions (p < 2.2 × 10⁻¹⁶; Fig. 7E). Together, the convergence of our filtered dREGs with both a reporter-based functional enhancer assay (Fig. 7C,D) and this independent regulatory-element compendium (Fig. 7E) supports the conclusion that at least a subset of the loci prioritized by our TE/sRNA-informed filtering correspond to genuine regulatory regions.

## Discussion

In this study, we show that moderate drought reprograms nascent transcription in rice not only at annotated genes but also at thousands of bidirectionally transcribed candidate regulatory loci, revealing a drought-responsive landscape of enhancer-like transcription initiation. Integrating PRO-seq with ATAC-seq, MethylC-seq, sRNA-seq, and Pore-C supports a model in which drought-responsive regulatory proteins engage accessible, locally hypomethylated cis-regulatory regions that can exhibit enhancer-like bidirectional initiation, and we show that some of these are coupled to target-gene output.

Genome-wide maps of accessible chromatin and active histone marks suggest that plants harbour tens of thousands of candidate enhancers, but only a subset have been functionally validated, and plant enhancers show substantial heterogeneity in location, chromatin signatures, and context dependence (Beernink et al., 2024). These features potentially distinguish plant enhancers from their animal counterparts, where canonical marks (H3K4me1/me3, H3K27ac) and pervasive bidirectional eRNA transcription provide more consistent identification criteria. Here, nascent transcription (PRO-seq) provides a complementary, activity-first view of regulatory logic under drought, because run-on assays directly measure engaged RNA polymerases and capture unstable, short-lived transcripts often missed by steady-state RNA profiling.

A recent comprehensive study using capped small-RNA sequencing (csRNA-seq) across plant species reported that unstable, vertebrate-like enhancer RNAs are rare in plants and that promoters, rather than distal elements, showed the strongest enhancer activity in STARR-seq assays (McDonald et al., 2024). This raises the question of whether bidirectional transcription at plant intergenic loci reflects genuine enhancer activity or instead marks unannotated promoters or other non-enhancer transcription. Several considerations apply to our results. First, PRO-seq and dREG measure engaged polymerase activity at initiation regions rather than classifying transcript stability, so the dREG sites we identify represent active bidirectional initiation events irrespective of downstream transcript fate. Second, our TE- and small-RNA-aware filtering specifically removes the intergenic transcription most likely to arise from heterochromatin or silencing-associated contexts. Third, and relatedly, the chromatin environment of our filtered sites (local accessibility, local hypomethylation, and absence of sRNA targeting) is more consistent with active regulatory elements than with unannotated promoters, which typically show broader stable-transcript accumulation and H3K4me3 enrichment. We acknowledge that some candidates may correspond to unannotated promoters or other features that do not conform to classical enhancer definitions, particularly given the emerging view that the promoter–enhancer boundary may be more fluid in plants than in animals (McDonald et al., 2024; Silver et al., 2024).

Despite the low number of vertebrate-like eRNAs reported by csRNA-seq, independent evidence supports the existence of bidirectionally transcribed enhancers in plant genomes. The MaizeCODE project identified thousands of distal enhancers expressing bidirectional noncoding RNAs in maize, some organized into clusters reminiscent of animal super-enhancers and enriched in regions under domestication selection (Cahn et al., 2024); about half of these overlapped dREG-identified regions from a previous GRO-seq analysis (Lozano et al., 2021), supporting the reliability of dREG in grasses. In wheat, enhancer transcription detected by pNET-seq and GRO-seq correlated more strongly with STARR-seq activity than chromatin marks such as H3K27ac or H3K9ac, suggesting nascent transcription may be a particularly informative predictor of enhancer function in crops (Xie et al., 2022). In Arabidopsis, dynamic enhancer transcription has been linked to immune-gene reprogramming during pattern-triggered immunity (Zhang et al., 2022). Together, these studies indicate that although plant eRNAs may differ from their vertebrate counterparts, bidirectional transcription initiation does mark a functionally relevant subset of plant regulatory elements.

The idea that transcription itself marks active enhancers is well established in animals, and the evidence in plants is now accumulating rapidly. The earlier gap partly reflected technical limitations — high-quality plant nascent-RNA datasets were rare, and methods optimized for mammalian regulatory architectures may not fully capture plant regulatory transcription (McDonald et al., 2024). As nascent-transcription approaches are increasingly applied across plant species, the field is converging on a more nuanced view in which enhancer-like transcription exists in plants but may be less pervasive, more condition- and spatially dependent, or structurally distinct from mammalian eRNA production. Our study is an early step toward establishing nascent transcription as a robust readout of enhancer activity in plants: the observation that moderate drought reprograms thousands of dREG-defined initiation regions, many intergenic, adds a stress-specific, mechanistic dimension to this emerging picture. The stress-inducible nature of many candidate sites is notable, because plant eRNA transcription may be more readily detected under conditions that actively engage regulatory networks than under steady-state developmental conditions.

A challenge in plants is that intergenic transcription is not necessarily synonymous with enhancer activity, because plant genomes contain large TE-rich intergenic regions whose chromatin is shaped by RdDM and small RNAs, producing signals that can confound enhancer predictions. Recent work shows that active regulatory landscapes can sit adjacent to, or derive from, transposon sequences, and that enhancer-like regions may carry distinct boundary signatures tied to small-RNA pathways (Li et al., 2015; Makarevitch et al., 2015; Zhao et al., 2018). Our TE/sRNA assessment is therefore informative beyond serving as a filter: the sharp transitions we observe, with rising CHH methylation and stronger small-RNA signal as TE overlap increases, recapitulate the expectation that canonical *cis*-regulatory activity is enriched in accessible, locally hypomethylated chromatin, whereas TE-associated transcription is embedded in silenced contexts; the filtered, low-TE candidates retained focal accessibility and local hypomethylation (Fig. 4C, D) (Cosby et al., 2019). Our conservative threshold (≤ 20% TE overlap, no sRNA overlap) prioritizes a higher-specificity subset of drought-responsive intergenic dREGs arising from regulatory activity. We also recognize that additional regulatory elements may reside in TE-associated space; TE-derived sequences are increasingly recognized as contributors of regulatory innovation, sometimes serving as raw material for new enhancers or harbouring co-opted TF binding sites (Chuong et al., 2017; Marand et al., 2017). Relaxing these thresholds, combined with functional validation, may uncover a broader set of drought-responsive regulatory elements within TE-rich domains. These data do not by themselves demonstrate TE exaptation, but they nominate a subset of young, TE-linked, drought-responsive dREGs as candidates for future tests of TE-derived regulatory co-option.

Within this high-confidence set, the aggregation of multi-omic signatures aligns with a growing consensus about what plant enhancers look like in practice: focal accessibility, local DNA-methylation depletion, and enrichment of regulatory activity at defined initiation sites (Zhu et al., 2015; Joly-Lopez et al., 2020; Beernink et al., 2024). Importantly, these features appear in a drought-responsive manner: upregulated intergenic dREGs gain nascent transcription and accessibility under drought, while downregulated sites show the opposite, consistent with enhancers as dynamic modules that integrate environmental signals rather than static sequence features. Because plant enhancer chromatin signatures vary by tissue, stage, and context, the convergence of initiation, accessibility, and methylation strengthens inference more than any single hallmark alone. We note that our study lacks drought histone-modification data commonly used to define enhancer states; while these would refine candidate classification, the combination of nascent transcription, chromatin accessibility, and methylation provides a complementary and, in some respects, more direct readout of regulatory activity, especially given that histone-mark–enhancer relationships differ between plants and animals (Yan et al., 2019).

Many functionally important plant regulatory elements reside in local regions near genes and are difficult to separate cleanly from promoter-proximal regulation (Ricci et al., 2019). Plant enhancers are reported in proximal, intergenic, and genic contexts, including gene-rich regions; although classically defined in animals as position- and orientation-independent, enhancer activity in plants (and increasingly in animals) can depend on genomic context, including relative position and sometimes orientation. Our finding that genes associated with proximal drought-responsive intergenic dREGs tend to exhibit higher nascent transcription, and that gene-level changes track the direction of nearby dREG regulation, fits a local regulatory-neighbourhood model. Our proximal dREGs, within 1 kb of transcription start or end sites, may therefore represent position-dependent regulatory elements whose drought-responsive transcription helps fine-tune nearby gene expression. The abundance of genic dREGs, including intronic and promoter-proximal enrichment, suggests drought-responsive initiation is distributed across multiple regulatory layers, including alternative promoters or intragenic regulatory modules.

Long-range regulation is a defining feature of many enhancers and is increasingly tractable in plants. High-resolution chromatin-interaction maps in Arabidopsis and crops suggest that long-range contacts can connect candidate regulatory elements to active genes, providing a three-dimensional framework for enhancer–promoter communication that complements linear genomic distance (Deng et al., 2023). Stress can reshape these contacts: 3D architecture responds to environmental perturbation in rice, including heat stress (Liang et al., 2021), and drought responses have been linked to dynamic, H3K9ac-marked 3D interaction networks associated with OsbZIP23, a key drought-responsive bZIP TF (Chang et al., 2024). That study reports a genome-wide disconnection of H3K9ac-marked contacts under drought, over 10, 000 loops lost and partially recovered upon re-watering, and the drought-specific contacts it characterizes are promoter–promoter interactions organized around “super-promoter regions” rather than distal enhancer–promoter loops. Filtered intergenic dREGs were not only enriched at gene-connected loop anchors, but the direction of their drought response predicted the direction of the looped gene’s transcriptional change, consistent with a subset of these elements acting as *bona fide* distal regulators rather than merely co-occurring with active chromatin. Because the Pore-C maps are not under drought, a key next step is to test whether drought alters loop strength or usage at the specific dREG–gene pairs nominated here, in line with stress-responsive 3D reorganization observed elsewhere.

Motif enrichment within activated intergenic dREGs provides testable hypotheses about the upstream regulators of drought-induced enhancer activation. Enrichment of motifs for TF families with known roles in drought and hormone signaling, including bZIP, NAC, MYB, and WRKY, supports a model in which drought-responsive TFs also act through enhancer-like elements to coordinate transcriptional reprogramming. In rice, co-enrichment of multiple stress-responsive TF motif families within the same dREG peaks suggests combinatorial regulation, a feature increasingly recognized at animal enhancers and demonstrated at plant enhancers through saturation mutagenesis of light-responsive elements (Jores et al., 2021).

The overlap of our filtered intergenic dREGs with published STARR-seq enhancers provides important support for the enhancer identity of a subset of our candidates: 284 STARR-seq enhancers overlap our filtered dREG peaks, indicating that some of our transcription-defined candidates possess autonomous enhancer activity in an independent assay. We note that STARR-seq measures enhancer potential outside of the native chromatin environment, and a lack of overlap does not imply a lack of endogenous regulatory function.

From a translational perspective, the drought-responsive enhancer-like elements identified here are candidate targets for crop improvement. Natural variation at cis-regulatory regions is a major contributor to phenotypic diversity and adaptive traits in crops, and noncoding regulatory variants underlie agronomically important QTLs in rice and other cereals (Ricci et al., 2019; Wang et al., 2022a; Wang and Bart, 2025). With CRISPR-based genome editing, the elements identified here could serve as rational targets for engineering drought resilience without altering protein-coding sequences (Rodríguez-Leal et al., 2017; Zhu et al., 2020). Candidates converging across multiple evidence types provide a ranked starting point for such efforts.

This study has limitations that bound our conclusions and point to the most informative next experiments. Our analyses derive from a single tissue (juvenile leaves), one developmental stage, and one drought severity. Enhancer landscapes are tissue- and condition-specific in both animals and plants (Andersson et al., 2014; Marand et al., 2021), and drought-responsive programs in other organs (e.g. roots) may involve different elements; stress intensity and duration also activate distinct pathways (Todaka et al., 2015), and our moderate treatment captures one point along that range. Applying the PRO-seq/dREG framework across tissues, stages, and stress regimes would enable a comprehensive atlas of condition-dependent enhancer-like transcription. In addition, dREG was trained on mammalian initiation signatures; although its utility in plants is supported by multiple studies, plant-trained models or newer tools could further improve detection. Finally, causal validation will require functional perturbation of prioritized loci through reporter assays, such as plant-adapted STARR-seq, and endogenous CRISPR editing, especially for elements nominated by multiple orthogonal signals.

In conclusion, our results support a working model in which drought reshapes nascent transcription at both genes and enhancer-like intergenic elements in rice. Moderate water deficit drives broad transcriptional reprogramming at annotated genes involved in defense, hormone signaling, and senescence, and simultaneously activates and represses thousands of dREG-defined transcription-initiation sites distributed across accessible, hypomethylated chromatin. TE-and sRNA-based filtering enriched for intergenic dREGs with enhancer-like chromatin and transcriptional hallmarks, and the activity of these filtered elements is associated with elevated gene expression, 3D contacts to target genes, and enrichment for stress-related TF motifs. We propose that several drought-responsive dREGs act as enhancer-like elements integrating chromatin context, TF binding, and 3D architecture to fine-tune stress-responsive transcriptional reprogramming in rice.

## Materials and Methods

### Plant cultivation and stress treatment

*Oryza sativa* ssp. japonica var. Azucena (PI 439017; tropical japonica) was cultivated in soil (PRO-MIX BX MYCORRHIZAE, 10381RG; Premier Tech, Canada) supplemented with Jack’s Professional General Purpose fertilizer (20-20-20; 384.7 mg per kg soil) and FeSO₄ (67.2 mg per kg soil), under controlled conditions in growth chambers (Conviron GEN1000; Conviron, Canada) with a 14 h/10 h day–night cycle, 28 °C/20 °C day/night temperature, 300–500 µmol quanta m⁻² s⁻¹, and 80% relative humidity. Well-watered conditions were defined by a soil-water content of 2.5 g H₂O per g soil, and drought stress by 1.25 g H₂O per g soil. Germinated seeds were planted in soil and kept under well-watered conditions for the first five days to promote seedling establishment. Drought was induced on day five by withholding water until the soil-water content reached the drought level, which was then maintained until tissue harvest. Seedlings were grown for a total of 15 days before harvesting.

### PRO-seq

The PRO-seq protocol was based on (Hetzel et al., 2016), and (Mahat et al., 2016). Nuclei were isolated from 15-day-old seedling-derived tissue homogenized in grinding buffer (300 mM sucrose, 20 mM Tris-HCl pH 8.0, 5 mM KCl, 5 mM Triton X-100, 35% glycerol, 5 mM β-mercaptoethanol, protease-inhibitor cocktail, RNase inhibitor, DEPC-H₂O) using a TissueRuptor II. The homogenate was filtered through two 250-µm nylon meshes (Fisher Scientific, NC0148095), one layer of Miracloth (Millipore Sigma, 475855), and a 40-µm cell strainer (Corning). Nuclei were pelleted by centrifugation (10 min, 5,000 × g, 4 °C), homogenized in cold grinding buffer with a loose Kimble dounce and resuspended in storage buffer (10 mM Tris -HCl pH 8.0, 25% glycerol, 5 mM MgCl₂, 0.1 mM EDTA, 5 mM DTT, DEPC-H₂O). Nuclei were stained and counted on a BD FACSJazz cell sorter and diluted to 5–10 × 10⁶ nuclei per sample. Sequencing libraries were prepared as described in Mahat et al. 2016 with a four-biotin run-on and sequenced as 2 × 100 bp on an Illumina NovaSeq 6000 (Génome Québec, Québec, Canada).

### ATAC-seq

ATAC-seq was performed according to Buenrostro et al. 2015 and Zhang & Jiang 2015, from frozen plant tissue. Flash-frozen tissue (0.2–0.3 g) was ground to a fine powder and dissolved in ice-cold nuclei-isolation buffer (NIB; 10 mM Tris-HCl, 80 mM KCl, 10 mM EDTA, 1 mM spermidine, 1 mM spermine, 0.15% β-mercaptoethanol, 0.5 M sucrose, pH 9.5), filtered through Miracloth (Millipore Sigma, 475855), and pelleted by centrifugation (1,100 × g, 10 min, 4 °C). The pellet was washed twice in 10 ml nuclei-washing buffer (NIB with 0.5% Triton X-100; Sigma, T8787) and re-pelleted, equilibrated in 10 ml nuclei-digestion buffer (10 mM Tris-HCl, 10 mM NaCl, 3 mM MgCl₂, pH 7.4), and resuspended in resuspension buffer (20 mM Tris-HCl, 10 mM NaCl, 3 mM MgCl₂, pH 7.4). Nuclei were stained with propidium iodide, counted on a BD FACSJazz cell sorter, and pelleted (1,100 × g, 10 min, 4 °C). A total of 50,000 nuclei were resuspended in a transposition reaction mix (Tagment DNA buffer, Tagment DNA Enzyme TDE1; Illumina, 20034210) and incubated at 37 °C for 30 min with gentle shaking. Samples were purified with the QIAGEN MinElute PCR Purification Kit (28004) and eluted in 10 µl elution buffer, amplified and tagged with custom sequencing primers, purified a second time with the same kit, and size-selected with AMPure XP beads (Beckman Coulter, A63880). Libraries were sequenced as 2 x 100 bp on an Illumina NovaSeq 6000.

Sequenced reads were quality-checked with FastQC and 3′-trimmed to remove adapters and low-quality bases (Q > 20) with Trimmomatic v0.39 (Bolger et al., 2014); HEADCROP:10, MINLEN:16, SLIDINGWINDOW:5:20, then aligned to the *O. sativa* japonica Azucena reference genome (Ensembl Plants release 57) with Bowtie2 v2.4.4 (Langmead and Salzberg, 2012); --very-sensitive -X 1000. The reference included the chloroplast sequence (GenBank GU592207.1) so that chloroplast-derived reads could be captured and subsequently removed with Samtools v1.17 (Danecek et al., 2021). Duplicates were removed with Picard MarkDuplicates v2.26.3, and reads were shifted +4 bp on the plus strand and −5 bp on the minus strand to account for transposase binding, using deepTools v3.5.4 (Ramírez et al., 2016). Peaks were called with MACS2 v2.2.8 (Zhang et al., 2008) genome size and parameters recommended for ATAC-seq (-q 0.01 -g 379627553 --nomodel --shift 37 --extsize 73); narrow peaks were called, as these correspond to the smaller accessible regions that typically represent transcription-factor binding sites and regulatory elements. FRiP scores were computed with featureCounts (subread v2.0.6). An informed signal cutoff of 350 was applied to retain high-confidence ATAC peaks: at this threshold we observed a gradual enrichment of ATAC peaks at dREG sites together with a decrease in peak coverage in the 500 bp upstream of the TSS (Table S6).

### MethylC-seq

MethylC-seq was performed as described in (Urich et al., 2015). Genomic DNA was extracted from leaf tissue of 15-day-old seedlings with the QIAGEN DNeasy Plant Mini Kit and fragmented to ∼200 bp with a Bioruptor Plus sonicator (Diagenode). Fragments were size-selected with AMPure XP beads (Beckman Coulter). End repair was performed with the End-It DNA End-Repair Kit (VWR International, ER81050), followed by two AMPure XP purifications. Adapter-ligated gDNA was then bisulfite-converted with the EZ DNA Methylation-Lightning Kit (Zymo Research, D5030) according to the manufacturer’s instructions and purified with a final AMPure XP step (ratio 1:1). Libraries were sequenced as 2 x 100 bp reads on an Illumina NovaSeq 6000.

Sequencing data were processed with cutadapt v2.10 (Martin, 2011) and methylpy v1.4.6 (Schultz et al., 2015). methylpy built a bisulfite-converted reference index and aligned the paired-end reads. Per-sample bisulfite conversion rates (> 99.3% for all libraries) were estimated from the unmethylated chloroplast control, and genome-wide weighted methylation levels were computed per sequence context (CG, CHG, CHH) from the methylpy allc output (Table S7). The allc files were converted to bigWig format for downstream analyses.

### Bioinformatic analysis of nascent transcription and dREG calling

PRO-seq reads were processed with the proseq2.0 pipeline (github.com/Danko-Lab/proseq2.0), which pre-processes the reads, aligns them to the reference genome, and produces strand-specific bigWig files for peak calling; alignments were filtered to a minimum mapping quality of 20 (Samtools, -q 20). Replicate reproducibility was assessed by pairwise correlation of PRO-seq read counts across biological replicates in gene bodies and in promoter-proximal windows, and visualized with ggplot2 (geom_hex, geom_abline) and ggpubr (stat_cor) (Supplementary Figs. S1 and S2). Bidirectionally transcribed regulatory loci were called with dREG (Danko et al., 2015; Wang et al., 2019) independently for each biological replicate. Consensus peaks were retained when peaks from two or more replicates of a condition overlapped by ≥ 1 bp after extending coordinates by ± 100 bp, yielding 46,101 well-watered and 39,770 drought consensus peaks (Tables S2 and S9). The promoter pause index was computed per gene (> 5 kb) as the ratio of PRO-seq signal in a promoter-proximal window to the gene body, following the analysis scripts of Judd et al. 2020 (github.com/JAJ256/qPRO), and per-gene differences between conditions were tested with a Wilcoxon test (Benjamini–Hochberg correction). The intronic-to-exonic read ratio was computed from PRO-seq signal over annotated introns versus exons.

### Differential analysis of genes and dREGs

Differentially expressed genes and dREGs were identified with DESeq2 within the tfTarget framework (Chu et al. 2018) from PRO-seq gene-body and dREG-peak counts, respectively. DEGs were defined at padj ≤ 0.05 and |log2FC| ≥ 1; DE dREGs at padj ≤ 0.05, with a more stringent |log2FC| ≥ 0.5 applied only for directionally stratified analyses. GO enrichment used the clusterProfiler R package v4.14.6 (Yu et al., 2012; Wu et al., 2021) against a genome background.

### Genomic annotation, intron overlap, and TE/sRNA filtering

dREG peaks were annotated to genomic features with the annotatr R package, using a panel of BED files defining distal intergenic, upstream (500 bp and 500–1,000 bp), promoter-proximal (TSS ± 100 bp and TSS + 500 bp), genic (TSS-to-CDS, exonic and CDS-intronic) and downstream (500 bp and 500–1,000 bp) regions derived from the Azucena gene models, together with transposable-element and small-RNA annotations. Every annotated region overlapping a peak by at least 1 bp was recorded; peaks overlapping any genic category were classified as genic, and peaks overlapping only distal, upstream or downstream windows as intergenic. Intersections with annotated introns, transposable elements, and intergenic small-RNA loci were computed with bedtools v2.31.0. Intergenic DE dREGs with TE overlap ≤ 20% and no intergenic sRNA overlap were retained as the filtered enhancer-like set (2,428 peaks).

### Young TE enrichment analysis

To test whether TE-linked sequence was retained among drought-responsive enhancer-like candidates after TE/sRNA filtering, we intersected the 841 upregulated filtered intergenic dREGs with annotated transposable elements using bedtools v2.31.0 intersect. The foreground consisted of upregulated filtered intergenic dREGs defined by padj ≤ 0.05, log2FC ≥ 0.5, TE overlap ≤ 20%, and no intergenic sRNA overlap. TE age was estimated from sequence divergence to the corresponding TE-family consensus, with lower-divergence copies considered younger. Young TE overlaps were tested for enrichment by major TE class using two-sided Fisher’s exact tests in R, with Benjamini–Hochberg correction for multiple testing. The background consisted of filtered intergenic dREGs passing the same TE/sRNA criteria. Odds ratios and adjusted P values were visualized as a forest plot.

### Chromatin meta-profiles and pause-index figures

Within-and-around meta-profiles and heatmaps of PRO-seq, ATAC-seq, and MethylC-seq signal within ± 1 kb of filtered dREG peaks were generated with deepTools (computeMatrix/plotProfile/plotHeatmap), stratified by direction of regulation and by condition.

### Pore-C loop enrichment

Filtered intergenic dREGs were intersected with Pore-C loop anchors from Azucena leaves (Goliasse et al., 2025). Enrichment of dREGs at gene-connected anchors was tested by a bootstrap permutation (1,000 iterations) against control regions matched on length and distance-to-nearest-gene, restricted to the ascertainable intergenic space; significance was assessed as the fraction of permutations reaching the observed count (observed 3.19-fold, p < 0.001).

### Motif enrichment

*De novo* and known-motif enrichment on filtered upregulated intergenic dREG peaks (n = 841; 6–8 bp) was performed with HOMER (findMotifsGenome.pl) against a genome-matched background, separately for proximal and distal subsets.

### STARR-seq comparison

A published rice protoplast STARR-seq enhancer set (9,642 elements; (Sun et al., 2019), Nipponbare coordinates) was lifted to the Azucena assembly with liftOver (retaining 8,680 elements) and intersected with the TE/sRNA-filtered dREG catalogue (16,828 peaks) within a ± 200 bp window around each dREG summit. Bidirectionality of PRO-seq signal at overlapping regions was scored per strand and compared between conditions with a paired Wilcoxon test.

### Statistics and data visualization

Statistical tests are indicated with each result; multiple testing was controlled by the Benjamini–Hochberg procedure. The following software was used: Trimmomatic v0.39, Bowtie2 v2.4.4, Samtools v1.17, Picard v2.26.3, MACS2 v2.2.8, subread/featureCounts v2.0.6, cutadapt v2.10, methylpy v1.4.6, bedtools v2.31.0, deepTools v3.5.4 (computeMatrix, plotProfile and plotHeatmap), HOMER v5.1 (findMotifsGenome.pl; -mset plants -len 6,7,8), UCSC liftOver (-minMatch 0.8), tfTarget (github.com/Danko-Lab/tfTarget), and the R packages DESeq2 v1.46.0, clusterProfiler v4.14.6, EnhancedVolcano v1.24.0, annotatr v1.39.0, ggplot2 v4.0.3 and ggpubr.

### Data availability

Raw and processed PRO-seq, ATAC-seq, and MethylC-seq data generated in this study are in the process of being deposited in the NCBI Gene Expression Omnibus (GEO) and the Sequence Read Archive (SRA). Accession numbers, together with the analysis code, will be added to this manuscript and made publicly available prior to peer-reviewed publication. The two principal processed datasets underlying the main conclusions, the complete list of differentially expressed genes (Table S4) and the consensus dREG peak catalogue (Table S9), are provided with this preprint.

## Author contributions

FOK and ZJL designed the study; FOK, AJ, and ZJL collected data; FOK, FR, MG, JG, and ZJL analysed data; AJ assisted with analysis; FOK and ZJL drafted the manuscript with significant input from FR. All authors read and approved the final manuscript.

## Acknowledgements

We thank Geneviève Bourret and Grégoire Bonnamour at the CERMO-FC for sequencing and FACS support. This research was enabled in part by Calcul Québec and the Digital Research Alliance of Canada. We thank Genome Québec and the Centre de génomique CHU de Québec-Université Laval for the sequencing.

## Supplementary material

Supplementary Figures S1–S11 and Supplementary Tables S1–S8 are provided in the Supplementary Information file. Supplementary Tables S4 and S9 are large data tables and are provided as separate machine-readable files.

**Supplementary Table S4. Differentially expressed genes under drought.** Complete DESeq2 output for the 2,844 genes differentially transcribed between drought and well-watered conditions in the PRO-seq data (padj ≤ 0.05, |log₂FC| ≥ 1), computed from gene-body counts. Columns: *featureID* (Azucena gene identifier); *baseMean* (mean normalized count across all libraries); *log2FoldChange* (log₂ fold-change, drought versus well-watered; positive values indicate higher nascent transcription under drought); *lfcSE* (standard error of the log₂ fold-change); *pvalue* (Wald-test p-value); and *padj* (Benjamini–Hochberg adjusted p-value).

**Supplementary Table S9. Consensus dREG peak catalogue.** Genomic coordinates of all consensus dREG peaks called from the PRO-seq data, comprising 46,101 peaks in the well-watered condition and 39,770 peaks in the drought condition. Peaks were called with dREG independently for each biological replicate and retained when detected in at least two of three replicates within a condition (see Methods). BED-format columns: chromosome; peak start; peak end; dREG score; peak summit position; peak identifier (indicating the replicate of origin); and condition (control = well-watered, or drought).

