## Supplemental Figures and Tables for "Drought reshapes enhancer-like nascent transcription and gene regulation in *Oryza sativa*"

#### Supplementary Figures

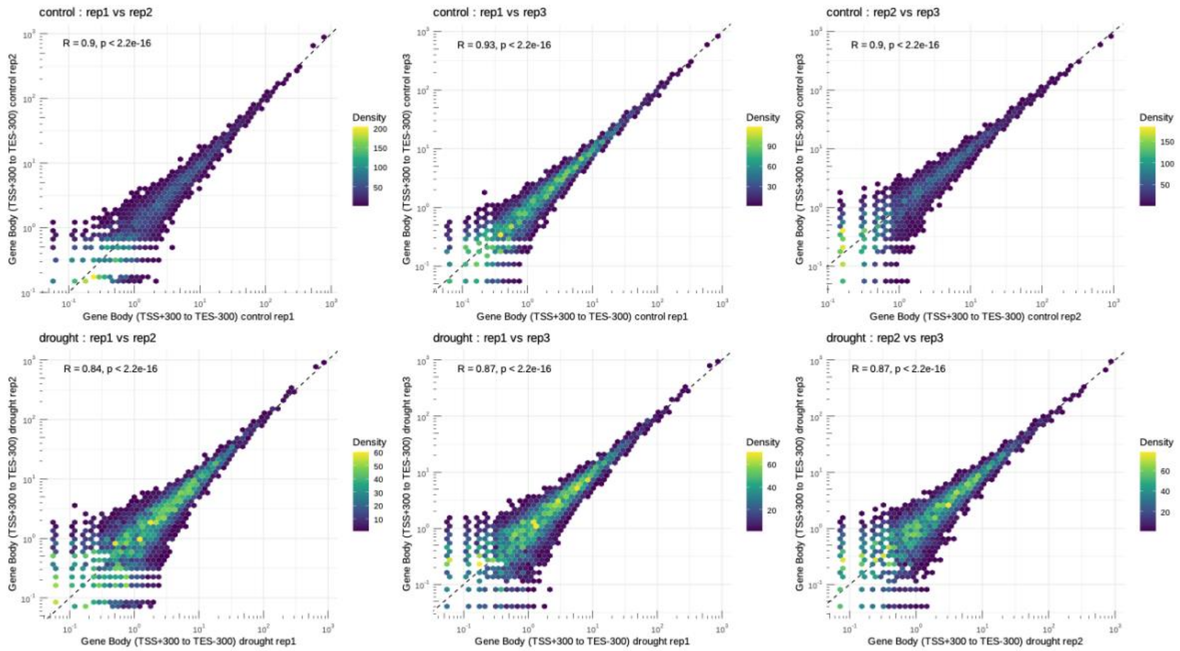


**Supplementary Figure S1. PRO-seq replicate correlation in gene bodies.** Pairwise correlation of PRO-seq read counts between biological replicates across gene-body regions (TSS + 300 bp to TES − 300 bp), for well-watered (top row) and drought (bottom row) conditions. Hexbin density plots are shown with the Pearson correlation coefficient (R) and p-value; the dashed line indicates identity.

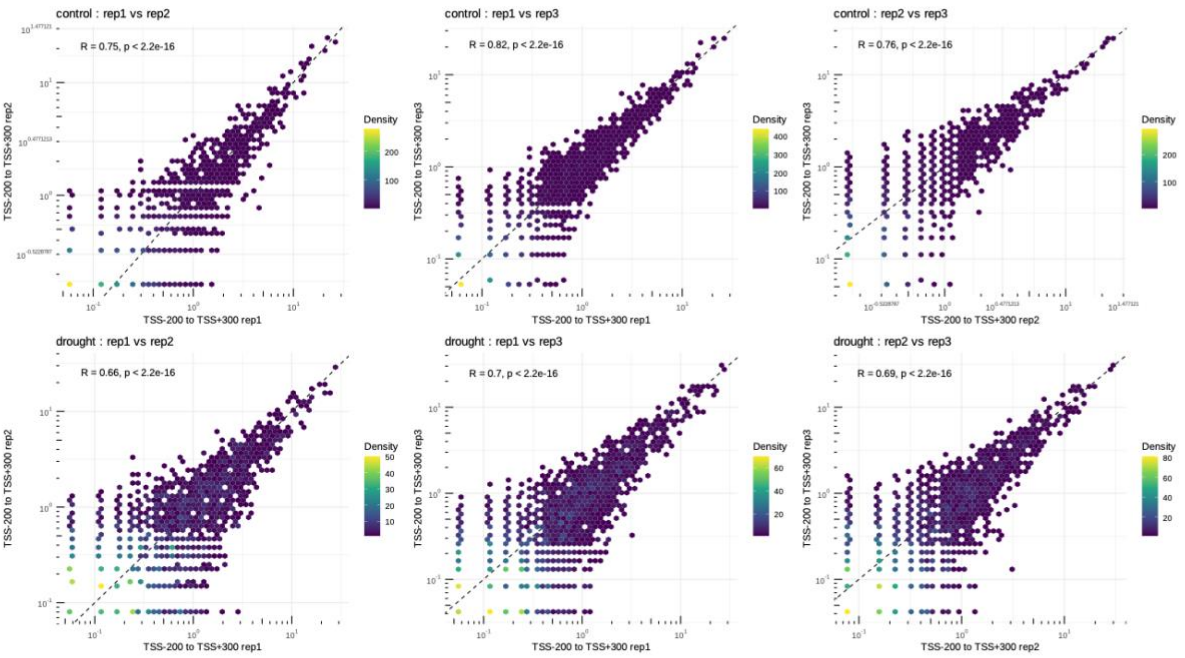


**Supplementary Figure S2. PRO-seq replicate correlation in promoter-proximal regions.** As in Supplementary Fig. S1, but for promoter-proximal windows (TSS − 200 bp to TSS + 300 bp).

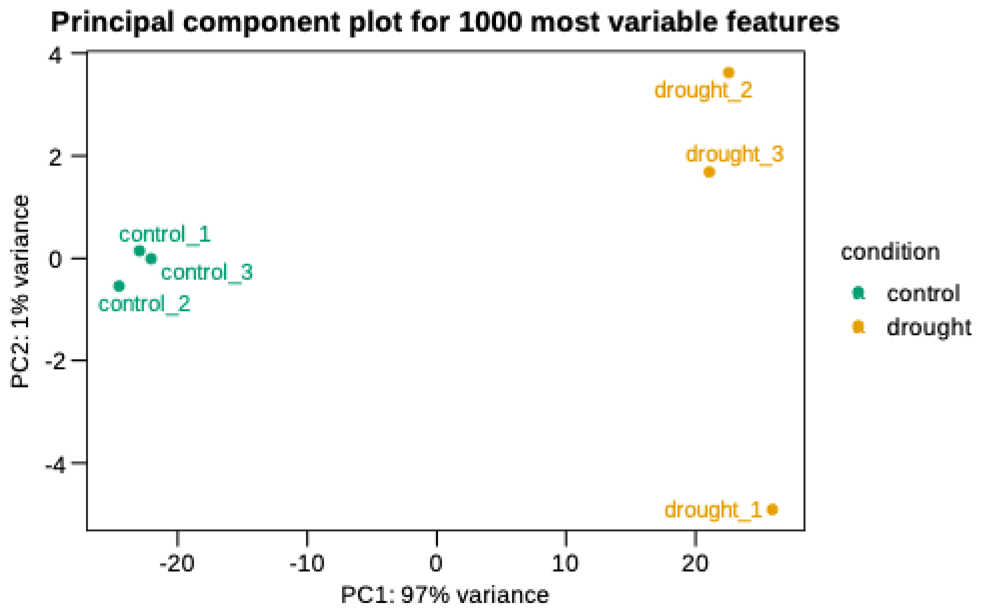


**Supplementary Figure S3. Principal-component analysis of PRO-seq replicates.** PCA of PRO-seq signal across the most variable features. Well-watered and drought samples separate along PC1 (97% of the variance), and biological replicates cluster tightly within each condition.

A

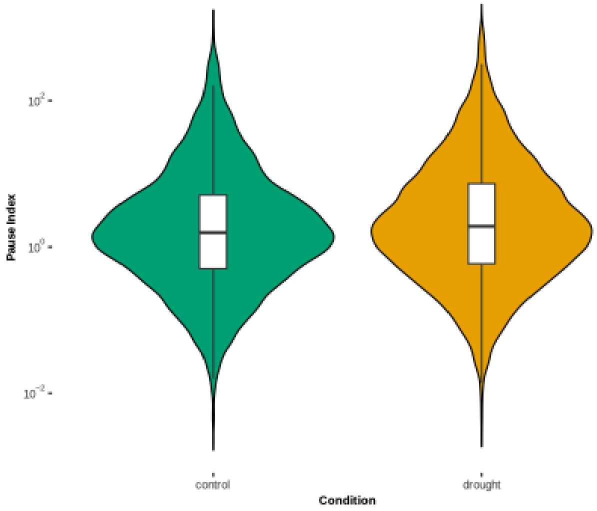


B

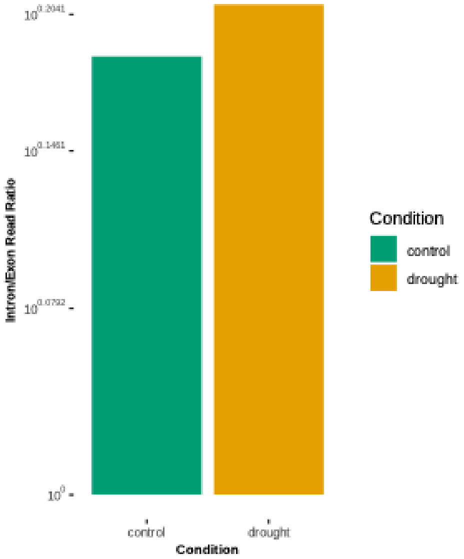


**Supplementary Figure S4. PRO-seq-based metrics of nascent transcription.** (A) Distribution of promoter-proximal pause indices for annotated genes under well-watered and drought conditions. The pause index was calculated as the ratio of PRO-seq signal in a promoter-proximal window to the signal across the downstream gene body for each gene. (B) Intron-to-exon read ratio for the PRO-seq libraries, calculated from reads mapping to intronic versus exonic regions of annotated genes.

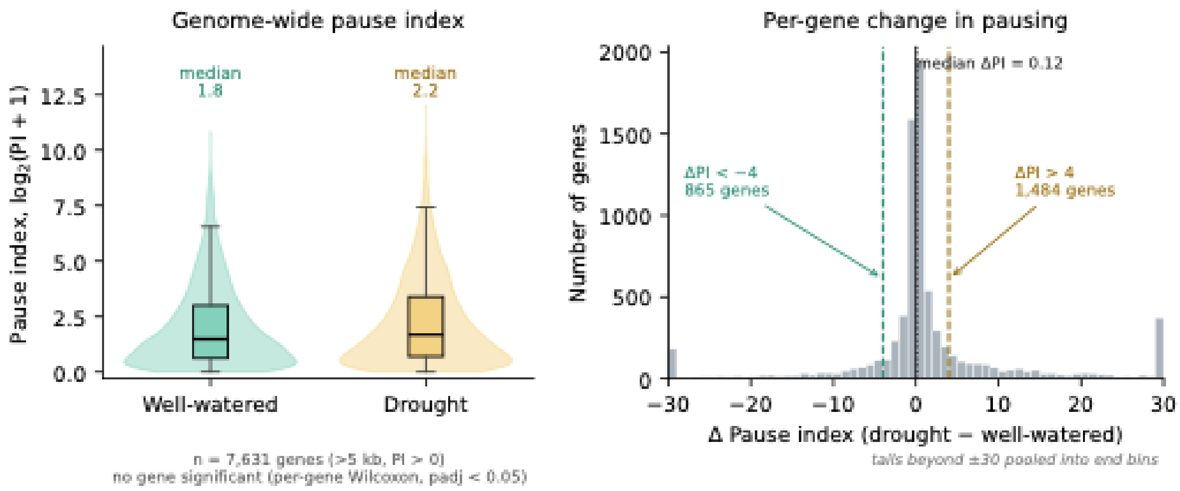


**Supplementary Figure S5. Pause-index differential analysis.** (A) Left: genome-wide per-gene pause index (PI) in well-watered and drought conditions, shown as $\log_2$(PI + 1), for the 7,631 genes longer than 5 kb passing PI > 0 filtering (median PI 1.8 versus 2.2; no individual gene significant by per-gene Wilcoxon test, Benjamini–Hochberg padj < 0.05). Right: per-gene change in pause index (ΔPI = drought − well-watered), centred near zero (median ΔPI = 0.12), with 1,484 genes at ΔPI > 4 and 865 genes at ΔPI < −4 (dashed lines); values beyond ±30 are pooled into the end bins. (B) Pause index in well-watered and drought conditions for the top 20 genes ranked by $\log_2$ fold-change among genes with ΔPI > 4, each shown on an independent y-axis (boxes = three biological replicates per condition).

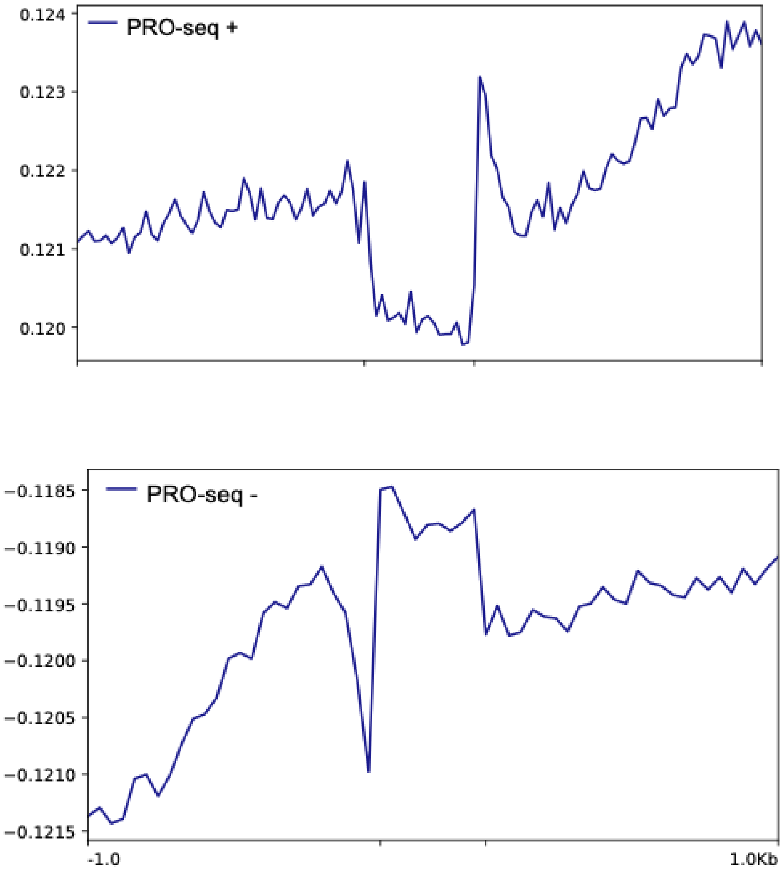


**Supplementary Figure S6. Divergent initiation at intron-overlapping dREG peaks.** Meta-profile of normalized PRO-seq signal centred on dREG peaks that overlap annotated introns, shown separately for plus- and minus-strand signal, confirming bidirectional transcription initiation within intronic sequences.

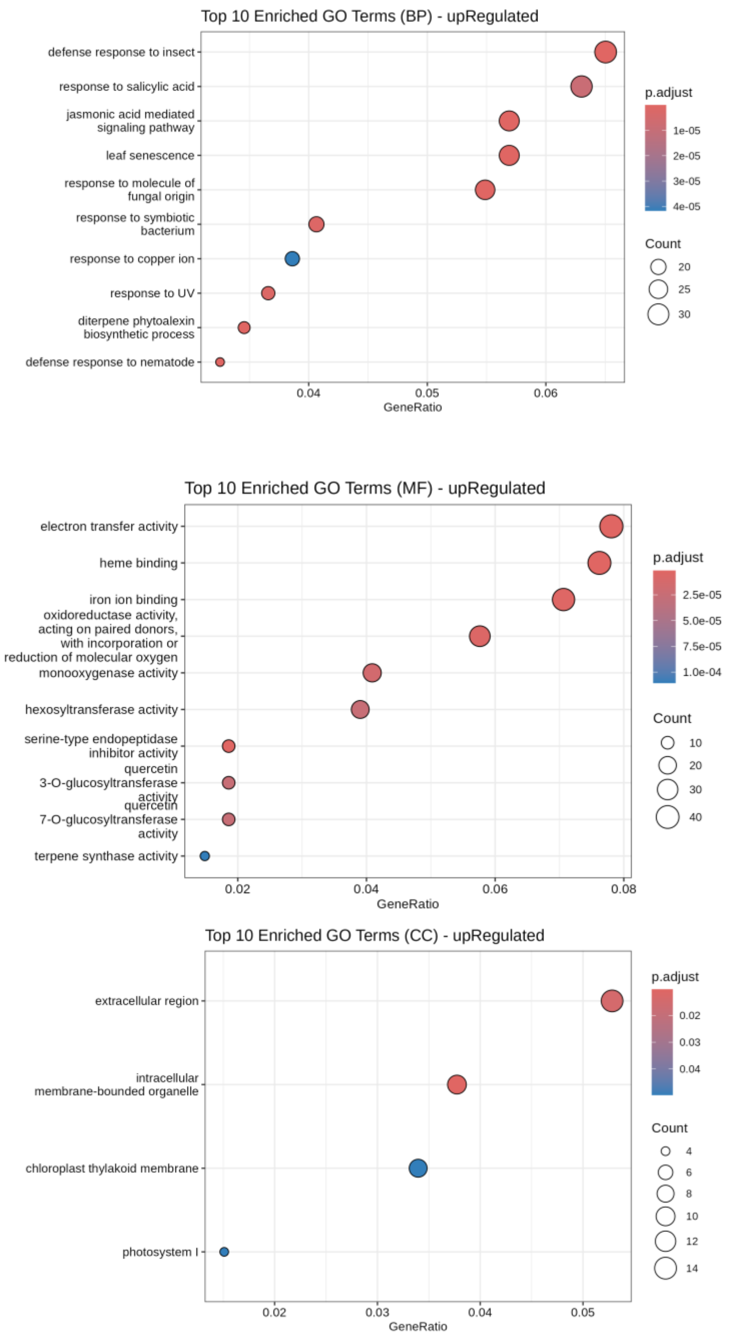


**Supplementary Figure S7. Gene Ontology enrichment for drought-upregulated genes.** GO enrichment (Biological Process, Molecular Function, Cellular Component) for genes upregulated under drought in the PRO-seq data (padj ≤ 0.05, $\log_2FC \geq 1$), using the full annotated gene set as background. Dot size indicates gene count; colour indicates the adjusted p-value.

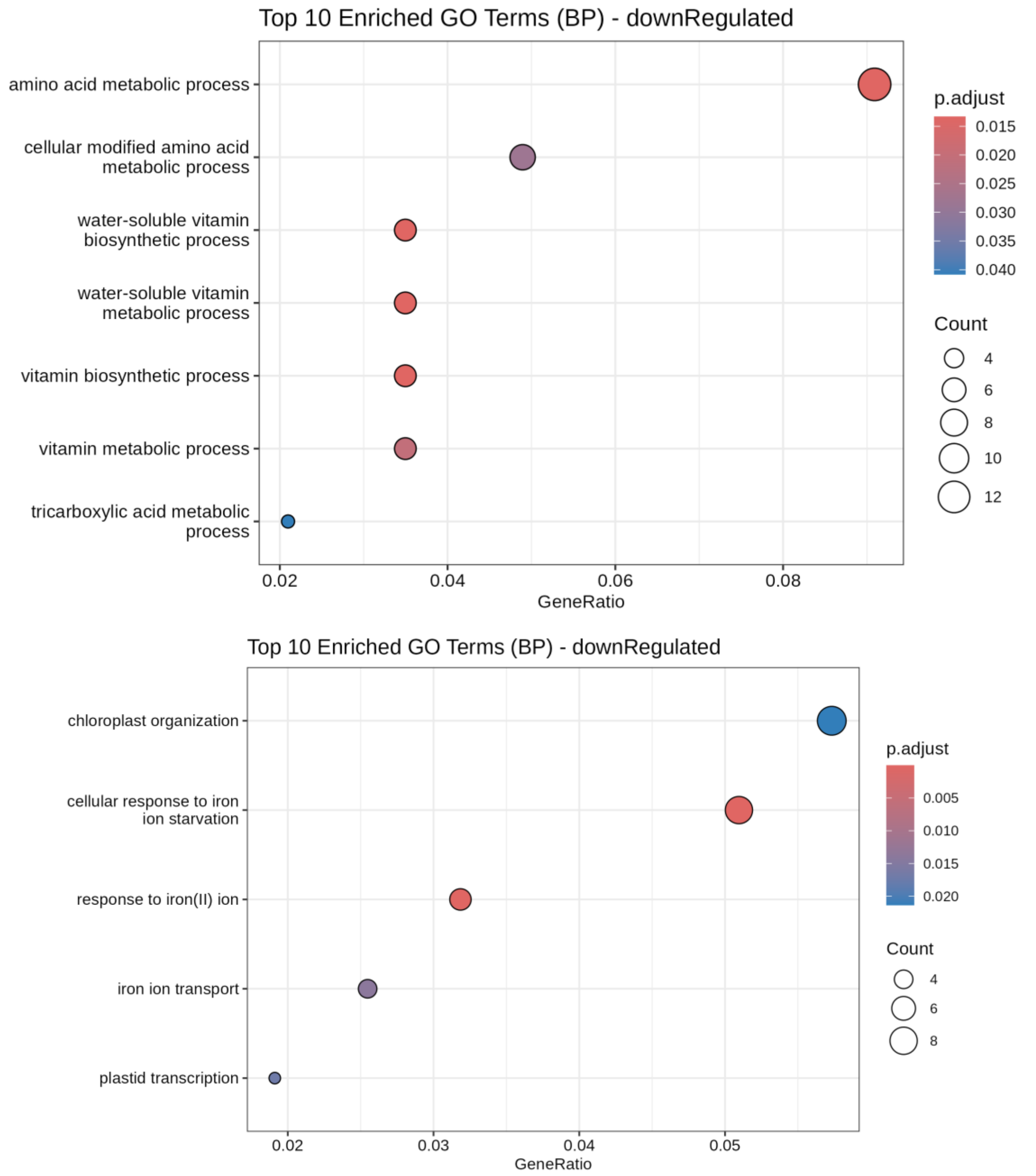


**Supplementary Figure S8. Gene Ontology enrichment for drought-downregulated genes.** As in Supplementary Fig. S7, for genes downregulated under drought (padj ≤ 0.05, $log_2FC \leq -1$).

**A**

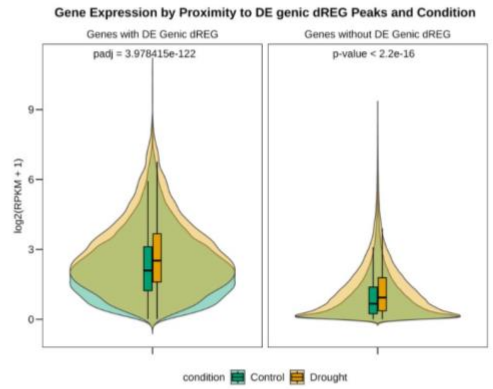


**B**

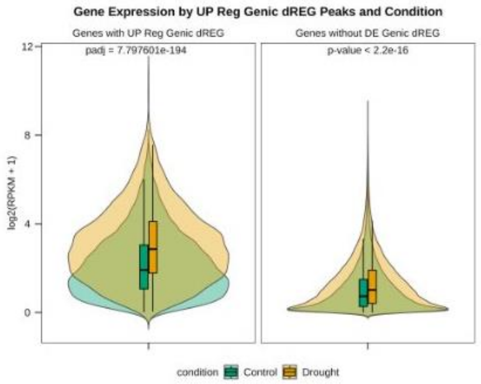


**C**

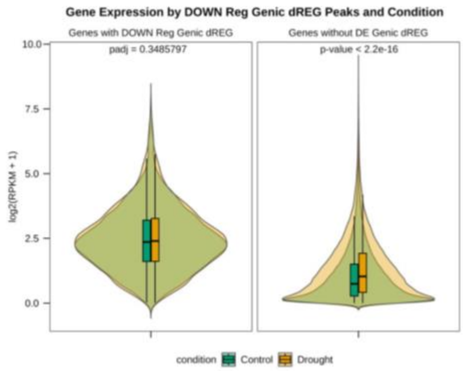


**Supplementary Figure S9. Gene expression proximal to genic differentially expressed dREG peaks.** Nascent transcription (normalized $log_2$ counts) of genes with versus without a differentially expressed genic dREG, in well-watered and drought conditions. (A) All genic DE dREGs. (B) Genes associated with upregulated genic dREGs (padj = $7.80 \times 10^{-194}$). (C) Genes associated with downregulated genic dREGs (padj = 0.35).

Known motifs

| Rank | Motif | Name | P-value | log P-pvalue | q-value (Benjamini) | # Target Sequences with Motif | % of Targets Sequences with Motif | # Background Sequences with Motif | % of Background Sequences with Motif | Motif File | SVG |
|---|---|---|---|---|---|---|---|---|---|---|---|
| 1 | | REM19(REM)/colamp-REM19-DAP-Seq(GSE60143)/Homer | 1e-9 | -2.138e+01 | 0.0000 | 231.0 | 27.47% | 8961.5 | 18.78% | motif file (matrix) | svg |
| 2 | | SPL5(SBP)/colamp-SPL5-DAP-Seq(GSE60143)/Homer | 1e-4 | -1.047e+01 | 0.0071 | 136.0 | 16.17% | 5472.6 | 11.47% | motif file (matrix) | svg |
| 3 | | SPCH(bHLH)/Seedling-SPCH-ChIP-Seq(GSE57497)/Homer | 1e-4 | -1.045e+01 | 0.0071 | 153.0 | 18.19% | 6311.5 | 13.23% | motif file (matrix) | svg |
| 4 | | NAM(NAC)/col-NAM-DAP-Seq(GSE60143)/Homer | 1e-4 | -1.023e+01 | 0.0071 | 181.0 | 21.52% | 7744.4 | 16.23% | motif file (matrix) | svg |
| 5 | | STZ(C2H2)/colamp-STZ-DAP-Seq(GSE60143)/Homer | 1e-4 | -9.369e+00 | 0.0086 | 447.0 | 53.15% | 22243.7 | 46.62% | motif file (matrix) | svg |
| 6 | | ANAC047(NAC)/colamp-ANAC047-DAP-Seq(GSE60143)/Homer | 1e-4 | -9.331e+00 | 0.0086 | 117.0 | 13.91% | 4681.6 | 9.81% | motif file (matrix) | svg |
| 7 | | AT5G02460(C2C2dof)/col-AT5G02460-DAP-Seq(GSE60143)/Homer | 1e-3 | -9.071e+00 | 0.0086 | 293.0 | 34.84% | 13801.8 | 28.93% | motif file (matrix) | svg |
| 8 | | bHLH80(bHLH)/col-bHLH80-DAP-Seq(GSE60143)/Homer | 1e-3 | -7.864e+00 | 0.0241 | 87.0 | 10.34% | 3404.9 | 7.14% | motif file (matrix) | svg |
| 9 | | WRKY40(WRKY)/colamp-WRKY40-DAP-Seq(GSE60143)/Homer | 1e-2 | -6.469e+00 | 0.0865 | 65.0 | 7.73% | 2511.2 | 5.26% | motif file (matrix) | svg |
| 10 | | LBD23(LOBAS2)/colamp-LBD23-DAP-Seq(GSE60143)/Homer | 1e-2 | -6.315e+00 | 0.0908 | 124.0 | 14.74% | 5434.7 | 11.39% | motif file (matrix) | svg |
| 11 | | PIF5ox(bHLH)/Arabidopsis-PIF5ox-ChIP-Seq(GSE35062)/Homer | 1e-2 | -6.172e+00 | 0.0953 | 112.0 | 13.32% | 4850.2 | 10.16% | motif file (matrix) | svg |
| 12 | | SPL1(SBP)/colamp-SPL1-DAP-Seq(GSE60143)/Homer | 1e-2 | -5.921e+00 | 0.1122 | 284.0 | 33.77% | 13975.3 | 29.29% | motif file (matrix) | svg |
| 13 | | PIF4(bHLH)/Seedling-PIF4-ChIP-Seq(GSE35315)/Homer | 1e-2 | -5.809e+00 | 0.1158 | 131.0 | 15.58% | 5874.2 | 12.31% | motif file (matrix) | svg |
| 14 | | bHLH130(bHLH)/col-bHLH130-DAP-Seq(GSE60143)/Homer | 1e-2 | -5.714e+00 | 0.1184 | 70.0 | 8.32% | 2836.3 | 5.94% | motif file (matrix) | svg |
| 15 | | AT1G47655(C2C2dof)/colamp-AT1G47655-DAP-Seq(GSE60143)/Homer | 1e-2 | -5.647e+00 | 0.1184 | 410.0 | 48.75% | 21027.7 | 44.07% | motif file (matrix) | svg |
| 16 | | SPL13(SBP)/col-SPL13-DAP-Seq(GSE60143)/Homer | 1e-2 | -5.562e+00 | 0.1205 | 26.0 | 3.09% | 824.7 | 1.73% | motif file (matrix) | svg |
| 17 | | CDF3(C2C2dof)/colamp-CDF3-DAP-Seq(GSE60143)/Homer | 1e-2 | -5.552e+00 | 0.1205 | 212.0 | 25.21% | 10174.7 | 21.32% | motif file (matrix) | svg |
| 18 | | SPL15(SBP)/colamp-SPL15-DAP-Seq(GSE60143)/Homer | 1e-2 | -5.321e+00 | 0.1363 | 234.0 | 27.82% | 11408.5 | 23.91% | motif file (matrix) | svg |
| 19 | | At5g47390(MYBrelated)/col-At5g47390-DAP-Seq(GSE60143)/Homer | 1e-2 | -5.147e+00 | 0.1537 | 154.0 | 18.31% | 7190.3 | 15.07% | motif file (matrix) | svg |

*De novo* motifs

| Rank | Motif | P-value | log P-pvalue | % of Targets | % of Background | STD(Bg STD) | Best Match/Details | Motif File |
|---|---|---|---|---|---|---|---|---|
| 1 | | 1e-17 | -4.131e+01 | 18.19% | 8.59% | 57.1bp (67.4bp) | GATA20(C2C2gata)/colamp-GATA20-DAP-Seq(GSE60143)/Homer(0.744) More Information \| Similar Motifs Found | motif file (matrix) |
| 2 | | 1e-14 | -3.326e+01 | 17.12% | 8.64% | 53.4bp (63.3bp) | At3g60580(C2H2)/col-At3g60580-DAP-Seq(GSE60143)/Homer(0.614) More Information \| Similar Motifs Found | motif file (matrix) |
| 3 | | 1e-12 | -2.916e+01 | 12.72% | 5.95% | 55.0bp (60.3bp) | P(MYB)/Zea mays/AthaMap(0.611) More Information \| Similar Motifs Found | motif file (matrix) |
| 4 | | 1e-12 | -2.900e+01 | 39.83% | 28.19% | 54.8bp (62.4bp) | At3g60580(C2H2)/col-At3g60580-DAP-Seq(GSE60143)/Homer(0.695) More Information \| Similar Motifs Found | motif file (matrix) |
| 5 | | 1e-12 | -2.807e+01 | 9.63% | 3.98% | 53.0bp (61.0bp) | At5g47390(MYBrelated)/col-At5g47390-DAP-Seq(GSE60143)/Homer(0.704) More Information \| Similar Motifs Found | motif file (matrix) |
| 6 | | 1e-12 | -2.768e+01 | 11.41% | 5.19% | 57.5bp (58.5bp) | TCP2/MA1064.1/Jaspar(0.822) More Information \| Similar Motifs Found | motif file (matrix) |
| 7 * | | 1e-11 | -2.732e+01 | 7.97% | 3.00% | 53.8bp (60.9bp) | PIF4(bHLH)/Seedling-PIF4-ChIP-Seq(GSE35315)/Homer(0.758) More Information \| Similar Motifs Found | motif file (matrix) |
| 8 * | | 1e-11 | -2.713e+01 | 38.17% | 27.07% | 53.8bp (69.4bp) | REM19(REM)/colamp-REM19-DAP-Seq(GSE60143)/Homer(0.844) More Information \| Similar Motifs Found | motif file (matrix) |
| 9 * | | 1e-10 | -2.469e+01 | 7.49% | 2.90% | 55.8bp (60.5bp) | At5g04390(C2H2)/col200-At5g04390-DAP-Seq(GSE60143)/Homer(0.772) More Information \| Similar Motifs Found | motif file (matrix) |
| 10 * | | 1e-9 | -2.277e+01 | 16.77% | 9.69% | 49.9bp (58.5bp) | NLP7(RWPRK)/col-NLP7-DAP-Seq(GSE60143)/Homer(0.740) More Information \| Similar Motifs Found | motif file (matrix) |
| 11 * | | 1e-9 | -2.209e+01 | 34.01% | 24.40% | 57.6bp (62.1bp) | Dof2/MA0020.1/Jaspar(0.712) More Information \| Similar Motifs Found | motif file (matrix) |
| 12 * | | 1e-9 | -2.186e+01 | 4.04% | 1.12% | 54.4bp (70.7bp) | TGA2(bZIP)/colamp-TGA2-DAP-Seq(GSE60143)/Homer(0.731) More Information \| Similar Motifs Found | motif file (matrix) |
| 13 * | | 1e-9 | -2.093e+01 | 7.49% | 3.19% | 51.7bp (66.4bp) | BHLH112/MA0961.1/Jaspar(0.695) More Information \| Similar Motifs Found | motif file (matrix) |
| 14 * | | 1e-8 | -1.967e+01 | 35.32% | 26.15% | 58.9bp (58.7bp) | Knotted(Homeobox)/Corn-KN1-ChIP-Seq(GSE39161)/Homer(0.702) More Information \| Similar Motifs Found | motif file (matrix) |
| 15 * | | 1e-7 | -1.633e+01 | 3.45% | 1.08% | 46.0bp (58.8bp) | GTL1(Trihelix)/colamp-GTL1-DAP-Seq(GSE60143)/Homer(0.661) More Information \| Similar Motifs Found | motif file (matrix) |
| 16 * | | 1e-7 | -1.618e+01 | 14.03% | 8.55% | 58.0bp (73.3bp) | SPL5(SBP)/colamp-SPL5-DAP-Seq(GSE60143)/Homer(0.958) More Information \| Similar Motifs Found | motif file (matrix) |
| 17 * | | 1e-7 | -1.614e+01 | 7.37% | 3.55% | 57.3bp (62.2bp) | MF0011.1_HMG_class/Jaspar(0.751) More Information \| Similar Motifs Found | motif file (matrix) |
| 18 * | | 1e-6 | -1.557e+01 | 2.50% | 0.63% | 35.9bp (57.5bp) | JGL(C2H2)/col-JGL-DAP-Seq(GSE60143)/Homer(0.699) More Information \| Similar Motifs Found | motif file (matrix) |
| 19 * | | 1e-6 | -1.489e+01 | 7.13% | 3.52% | 55.1bp (65.0bp) | MYB107(MYB)/col-MYB107-DAP-Seq(GSE60143)/Homer(0.651) More Information \| Similar Motifs Found | motif file (matrix) |

**Supplementary Figure S10. Full HOMER motif-enrichment results for drought-upregulated intergenic dREGs.** Known-motif and *de novo* motif logos recovered by HOMER from the 841 drought-upregulated filtered intergenic dREG peaks (6–8 bp) against a genome-matched background, with the best-matching transcription-factor family, enrichment p-value, and the percentage of target versus background sequences containing each motif.

Proximal

| Rank | Motif | P-value | log P-pvalue | % of Targets | % of Background | STD(Bg STD) | Best Match/Details | Motif File |
|---|---|---|---|---|---|---|---|---|
| 1 | GGTATCAT | 1e-14 | -3.378e+01 | 15.96% | 6.12% | 53.9bp (59.1bp) | CRC(C2C2YABBY)/col-CRC-DAP-Seq(GSE60143)/Homer(0.750)<br>More Information \| Similar Motifs Found | motif file (matrix) |
| 2 | AGTAGAAC | 1e-13 | -3.150e+01 | 12.50% | 4.27% | 55.4bp (62.2bp) | At3g60580(C2H2)/col-At3g60580-DAP-Seq(GSE60143)/Homer(0.809)<br>More Information \| Similar Motifs Found | motif file (matrix) |
| 3 | AAGGAAA | 1e-12 | -2.826e+01 | 28.65% | 16.12% | 54.3bp (57.5bp) | AT5G60130(ABI3VP1)/col-AT5G60130-DAP-Seq(GSE60143)/Homer(0.797)<br>More Information \| Similar Motifs Found | motif file (matrix) |
| 4 * | GAATCGAA | 1e-11 | -2.694e+01 | 8.65% | 2.53% | 49.7bp (62.8bp) | POL008.1_DCE_S_I/Jaspar(0.652)<br>More Information \| Similar Motifs Found | motif file (matrix) |
| 5 * | TGGTAGCG | 1e-11 | -2.684e+01 | 40.38% | 26.32% | 57.5bp (62.1bp) | CRF4(AP2EREBP)/colamp-CRF4-DAP-Seq(GSE60143)/Homer(0.622)<br>More Information \| Similar Motifs Found | motif file (matrix) |
| 6 * | ATCTGAC | 1e-11 | -2.627e+01 | 25.58% | 14.10% | 53.9bp (62.1bp) | AtGRF6(GRF)/col-AtGRF6-DAP-Seq(GSE60143)/Homer(0.775)<br>More Information \| Similar Motifs Found | motif file (matrix) |
| 7 * | CCACATGA | 1e-11 | -2.619e+01 | 10.58% | 3.66% | 49.4bp (60.9bp) | PIF4(bHLH)/Seedling-PIF4-ChIP-Seq(GSE35315)/Homer(0.781)<br>More Information \| Similar Motifs Found | motif file (matrix) |
| 8 * | CTTTTTTG | 1e-11 | -2.543e+01 | 44.81% | 30.68% | 53.4bp (67.1bp) | BBX31(Orphan)/col-BBX31-DAP-Seq(GSE60143)/Homer(0.774)<br>More Information \| Similar Motifs Found | motif file (matrix) |
| 9 * | TAGGCAGC | 1e-10 | -2.506e+01 | 8.65% | 2.68% | 61.5bp (63.8bp) | MYB116(MYB)/colamp-MYB116-DAP-Seq(GSE60143)/Homer(0.676)<br>More Information \| Similar Motifs Found | motif file (matrix) |
| 10 * | TCGTAC | 1e-10 | -2.456e+01 | 36.15% | 23.22% | 58.4bp (69.1bp) | SPL5(SBP)/colamp-SPL5-DAP-Seq(GSE60143)/Homer(0.838)<br>More Information \| Similar Motifs Found | motif file (matrix) |
| 11 * | CTCGTCAC | 1e-9 | -2.212e+01 | 16.15% | 7.82% | 54.1bp (66.5bp) | TGA2(bZIP)/colamp-TGA2-DAP-Seq(GSE60143)/Homer(0.790)<br>More Information \| Similar Motifs Found | motif file (matrix) |
| 12 * | ATATGCGC | 1e-9 | -2.149e+01 | 6.54% | 1.85% | 46.6bp (62.1bp) | PCF/Arabidopsis-Promoters/Homer(0.676)<br>More Information \| Similar Motifs Found | motif file (matrix) |
| 13 * | GTGCAGGA | 1e-8 | -1.896e+01 | 6.15% | 1.84% | 46.8bp (62.8bp) | POL011.1_XCPE1/Jaspar(0.636)<br>More Information \| Similar Motifs Found | motif file (matrix) |
| 14 * | AGCGATCG | 1e-8 | -1.883e+01 | 16.92% | 8.96% | 59.2bp (69.6bp) | GATA20(C2C2gata)/colamp-GATA20-DAP-Seq(GSE60143)/Homer(0.708)<br>More Information \| Similar Motifs Found | motif file (matrix) |
| 15 * | CCGGGAGC | 1e-8 | -1.856e+01 | 3.85% | 0.77% | 48.3bp (59.9bp) | POL013.1_MED-1/Jaspar(0.805)<br>More Information \| Similar Motifs Found | motif file (matrix) |
| 16 * | TGACCAGT | 1e-7 | -1.755e+01 | 26.54% | 16.92% | 55.2bp (61.5bp) | MYB81(MYB)/col-MYB81-DAP-Seq(GSE60143)/Homer(0.661)<br>More Information \| Similar Motifs Found | motif file (matrix) |
| 17 * | ATGCAATC | 1e-7 | -1.643e+01 | 13.27% | 6.73% | 56.3bp (60.0bp) | WUS1(Homeobox)/colamp-WUS1-DAP-Seq(GSE60143)/Homer(0.691)<br>More Information \| Similar Motifs Found | motif file (matrix) |
| 18 * | TAATCGTC | 1e-7 | -1.639e+01 | 2.69% | 0.42% | 33.1bp (60.1bp) | AtMYB77(MYB)/Arabidopsis thaliana/AthaMap(0.720)<br>More Information \| Similar Motifs Found | motif file (matrix) |
| 19 * | TTGCGCT | 1e-7 | -1.625e+01 | 16.15% | 8.90% | 52.6bp (65.4bp) | E2FA(E2FDP)/colamp-E2FA-DAP-Seq(GSE60143)/Homer(0.661)<br>More Information \| Similar Motifs Found | motif file (matrix) |
| 20 * | CCCCCCAC | 1e-6 | -1.574e+01 | 6.35% | 2.24% | 40.9bp (56.6bp) | ALFIN1(HD-PHD)/Medicago sativa/AthaMap(0.730)<br>More Information \| Similar Motifs Found | motif file (matrix) |
| 21 * | CCCCTGAT | 1e-6 | -1.556e+01 | 2.12% | 0.26% | 44.6bp (53.2bp) | RAV1(2)(AP2/EREBP)/Arabidopsis thaliana/AthaMap(0.720)<br>More Information \| Similar Motifs Found | motif file (matrix) |

Distal

| Rank | Motif | P-value | log P-pvalue | % of Targets | % of Background | STD(Bg STD) | Best Match/Details | Motif File |
|---|---|---|---|---|---|---|---|---|
| 1 | CTGGACCG | 1e-14 | -3.450e+01 | 6.85% | 0.66% | 54.3bp (63.8bp) | TCP16/MA0587.1/Jaspar(0.681)<br>More Information \| Similar Motifs Found | motif file (matrix) |
| 2 * | TGCCTATC | 1e-11 | -2.749e+01 | 13.71% | 3.96% | 52.2bp (59.6bp) | MYB116(MYB)/colamp-MYB116-DAP-Seq(GSE60143)/Homer(0.723)<br>More Information \| Similar Motifs Found | motif file (matrix) |
| 3 * | GGAAAAAA | 1e-9 | -2.276e+01 | 12.77% | 4.06% | 56.5bp (68.9bp) | REM19(REM)/colamp-REM19-DAP-Seq(GSE60143)/Homer(0.823)<br>More Information \| Similar Motifs Found | motif file (matrix) |
| 4 * | GGTTGCGT | 1e-9 | -2.154e+01 | 11.21% | 3.37% | 52.6bp (62.8bp) | NAC083/MA1043.1/Jaspar(0.823)<br>More Information \| Similar Motifs Found | motif file (matrix) |
| 5 * | TGGCGAC | 1e-9 | -2.094e+01 | 13.08% | 4.50% | 47.4bp (64.7bp) | ERF105(AP2EREBP)/colamp-ERF105-DAP-Seq(GSE60143)/Homer(0.725)<br>More Information \| Similar Motifs Found | motif file (matrix) |
| 6 * | TATGTAG | 1e-8 | -1.938e+01 | 22.74% | 11.25% | 54.7bp (59.5bp) | DDF1(AP2EREBP)/col-DDF1-DAP-Seq(GSE60143)/Homer(0.708)<br>More Information \| Similar Motifs Found | motif file (matrix) |
| 7 * | ATACCGCA | 1e-7 | -1.826e+01 | 6.23% | 1.29% | 57.9bp (61.8bp) | AT1G76870(Trihelix)/col-AT1G76870-DAP-Seq(GSE60143)/Homer(0.586)<br>More Information \| Similar Motifs Found | motif file (matrix) |
| 8 * | GCTCTATC | 1e-7 | -1.810e+01 | 14.02% | 5.55% | 44.6bp (59.9bp) | POL009.1_DCE_S_II/Jaspar(0.585)<br>More Information \| Similar Motifs Found | motif file (matrix) |
| 9 * | GGGTCAAC | 1e-7 | -1.730e+01 | 8.72% | 2.58% | 56.2bp (57.1bp) | WRKY63/MA1092.1/Jaspar(0.889)<br>More Information \| Similar Motifs Found | motif file (matrix) |
| 10 * | TCGTCGT | 1e-6 | -1.588e+01 | 17.76% | 8.57% | 52.4bp (70.1bp) | MGP(C2H2)/colamp-MGP-DAP-Seq(GSE60143)/Homer(0.718)<br>More Information \| Similar Motifs Found | motif file (matrix) |
| 11 * | ATCGAGA | 1e-6 | -1.409e+01 | 7.48% | 2.33% | 54.7bp (63.1bp) | GATA20(C2C2gata)/colamp-GATA20-DAP-Seq(GSE60143)/Homer(0.663)<br>More Information \| Similar Motifs Found | motif file (matrix) |
| 12 * | ACGAAAAG | 1e-5 | -1.312e+01 | 8.72% | 3.18% | 46.0bp (60.7bp) | CDF3(C2C2dof)/colamp-CDF3-DAP-Seq(GSE60143)/Homer(0.721)<br>More Information \| Similar Motifs Found | motif file (matrix) |
| 13 * | ACTAATTA | 1e-5 | -1.307e+01 | 5.92% | 1.64% | 50.7bp (58.3bp) | ATHB23(ZFHD)/col-ATHB23-DAP-Seq(GSE60143)/Homer(0.872)<br>More Information \| Similar Motifs Found | motif file (matrix) |
| 14 * | GGGCTGC | 1e-5 | -1.284e+01 | 5.61% | 1.51% | 46.9bp (60.9bp) | POL010.1_DCE_S_III/Jaspar(0.721)<br>More Information \| Similar Motifs Found | motif file (matrix) |
| 15 * | GCATGCT | 1e-5 | -1.270e+01 | 11.84% | 5.25% | 50.8bp (65.4bp) | FUS3/MA0565.1/Jaspar(0.923)<br>More Information \| Similar Motifs Found | motif file (matrix) |
| 16 * | AGTGTAC | 1e-5 | -1.232e+01 | 24.92% | 15.25% | 55.5bp (60.7bp) | SPL4/MA1058.1/Jaspar(0.719)<br>More Information \| Similar Motifs Found | motif file (matrix) |
| 17 * | ATGGAAGG | 1e-4 | -1.136e+01 | 4.67% | 1.21% | 52.9bp (58.4bp) | POL008.1_DCE_S_I/Jaspar(0.698)<br>More Information \| Similar Motifs Found | motif file (matrix) |
| 18 * | AACCGATA | 1e-4 | -1.013e+01 | 4.05% | 1.04% | 64.7bp (60.7bp) | MYB70(MYB)/col-MYB70-DAP-Seq(GSE60143)/Homer(0.836)<br>More Information \| Similar Motifs Found | motif file (matrix) |
| 19 * | AAAGACTT | 1e-4 | -9.664e+00 | 5.61% | 1.92% | 56.5bp (56.6bp) | DOF5.7/MA0984.1/Jaspar(0.626)<br>More Information \| Similar Motifs Found | motif file (matrix) |
| 20 * | ATACCCGC | 1e-3 | -8.498e+00 | 1.56% | 0.16% | 28.2bp (49.0bp) | DEL2(E2FDP)/col-DEL2-DAP-Seq(GSE60143)/Homer(0.643)<br>More Information \| Similar Motifs Found | motif file (matrix) |

**Supplementary Figure S11. Motif enrichment in proximal versus distal drought-upregulated intergenic dREGs.** HOMER de novo motif enrichment for drought-upregulated intergenic dREG peaks stratified by proximity to genes (proximal = within 1 kb of a TSS or TES; distal = all remaining peaks).

### Supplementary Tables

**Supplementary Table S1. PRO-seq sequencing and quality-control summary.** Per-library raw and trimmed read counts, mapping statistics, and duplication rates for the three well-watered and three drought PRO-seq replicates.

**Before trimming**

| Sample | % Dups | % GC | Average Read length | Mio. Seqs |
|---|---|---|---|---|
| Control 1 | 43,35 | 47,5 | 101 bp | 38,3 |
| Control 2 | 64,05 | 50 | 101 bp | 28,5 |
| Control 3 | 48 | 47 | 101 bp | 45,4 |
| Drought 1 | 50,7 | 50,5 | 101 bp | 47,1 |
| Drought 2 | 62,9 | 53 | 101 bp | 47,9 |
| Drought 3 | 46,4 | 51,5 | 101 bp | 64,5 |

**After trimming**

| Sample | % Dups | % GC | Average Read length | Mio. Seqs |
|---|---|---|---|---|
| Control 1 | 20,45 | 41,5 | 41 bp | 23,6 |
| Control 2 | 17,25 | 40,5 | 34 bp | 10,4 |
| Control 3 | 20,35 | 41,5 | 40 bp | 26,2 |
| Drought 1 | 37,65 | 44 | 39 bp | 31,8 |
| Drought 2 | 34,55 | 43 | 41 bp | 22,9 |
| Drought 3 | 28,8 | 43,5 | 43 bp | 42,5 |

**Supplementary Table S2. dREG peaks called per replicate.** Number of dREG peaks called in each PRO-seq replicate before and after removal of peaks overlapping the plastid genome (GenBank GU592207.1), and the number of consensus peaks retained per condition (present in ≥ 2 of 3 replicates).

| Samples | Total Peaks | GU592207.1 Peaks | Final Peaks |
|---|---|---|---|
| dREG_control_rep1.dREG.peak.score | 38731 | 27 | 38704 |
| dREG_control_rep2.dREG.peak.score | 17229 | 20 | 17209 |
| dREG_control_rep3.dREG.peak.score | 38469 | 29 | 38440 |
| dREG_drought_rep1.dREG.peak.score | 21259 | 23 | 21236 |
| dREG_drought_rep2.dREG.peak.score | 19098 | 27 | 19071 |
| dREG_drought_rep3.dREG.peak.score | 38576 | 27 | 38549 |

**Supplementary Table S3. Genomic annotation of dREG peaks.** Distribution of consensus dREG peaks across genomic features (genic and intergenic categories, promoter-proximal windows, exons and introns), including the number and percentage of peaks overlapping annotated introns by ≥ 1 bp and lying entirely within introns, for each condition.

| **Feature category** | **Well-watered** | | **Drought** | |
|---|---|---|---|---|
| | **Count** | **%** | **Count** | **%** |
| **Summary (per condition)** | | | | |
| All consensus dREG peaks | 46,101 | 100.0 | 39,770 | 100.0 |
| **Genomic feature distribution (Fig. 1D)** | | | | |
| Distal intergenic | 16,501 | 35.8 | 12,864 | 32.3 |
| Upstream 500–1,000 bp | 3,421 | 7.4 | 2,601 | 6.5 |
| Upstream ≤ 500 bp | 1,684 | 3.7 | 1,339 | 3.4 |
| Downstream ≤ 500 bp | 4,765 | 10.3 | 4,803 | 12.1 |
| Downstream 500–1,000 bp | 4,529 | 9.8 | 4,336 | 10.9 |
| Intergenic sRNA+ | 3,896 | 8.5 | 3,236 | 8.1 |
| TE+ | 239 | 0.5 | 199 | 0.5 |
| TSS ± 500 bp | 2,397 | 5.2 | 2,138 | 5.4 |
| Exon | 3,152 | 6.8 | 3,679 | 9.3 |
| CDS-intronic | 5,521 | 12.0 | 4,444 | 11.2 |
| TE_sRNA+ | 12 | 0.0 | 6 | 0.0 |
| Unclassified | 188 | 0.4 | 253 | 0.6 |
| **Overlap with annotated introns** | | | | |
| Overlapping annotated introns (≥ 1 bp) | 7,474 | 16.2 | 6,544 | 16.5 |
| Entirely intronic | 3,876 | 8.4 | 2,916 | 7.3 |

**Supplementary Table S5. Transposable-element and small-RNA overlap of intergenic differentially expressed dREG peaks.** Per-peak fraction of overlap with annotated transposable elements, TE class, and overlap with intergenic small-RNA loci; these values define the filtered enhancer-like set (≤ 20% TE overlap, no intergenic sRNA overlap).

| **region_class** | **total_unique peaks** | **Up regulate** | **Down regulate** | **Total regulate** | **sRNA** | **TE** | **sRNA_DE** | **group** | **total peaks** | **DE peaks** |
|---|---|---|---|---|---|---|---|---|---|---|
| **intergenic** | 4522 | 371 | 724 | 1095 | - | - | - | | | |
| **intergenic_TE** | 51987 | 5089 | 4329 | 9418 | - | 51987 | - | | | |
| **TE** | 438 | 52 | 48 | 100 | - | 438 | - | **intergenic** | **64081** | **11926** |
| **intergenic_sRNA** | 73 | 8 | 11 | 19 | 73 | - | 19 | | | |
| **intergenic_TE_sRNA** | 7061 | 831 | 463 | 1294 | 7061 | 7061 | 1294 | | | |
| **genic_intergenic** | 1021 | 109 | 185 | 294 | - | - | - | | | |
| **genic_intergenic_sRNA** | 16 | - | 1 | 1 | 16 | - | 1 | | | |
| **genic_intergenic_TE** | 5966 | 848 | 496 | 1344 | - | 5966 | - | | | |
| **genic_intergenic_TE_sRNA** | 778 | 95 | 49 | 144 | 778 | 778 | 144 | | | |
| **genic** | 3566 | 360 | 819 | 1179 | - | - | - | **genic** | **21349** | **5143** |
| **genic_sRNA** | 13 | 2 | 2 | 4 | 13 | - | 4 | | | |
| **genic_TE** | 9077 | 1277 | 748 | 2025 | - | 9077 | - | | | |
| **genic_TE_sRNA** | 894 | 103 | 43 | 146 | 894 | 894 | 146 | | | |
| **TE_sRNA** | 18 | 6 | - | 6 | 18 | 18 | 6 | | | |
| **unclassified** | 441 | 25 | 130 | 155 | - | - | - | **removed** | **441** | **155** |
| **All** | 85871 | 9176 | 8048 | 17224 | 8853 | 76219 | 1614 | **total** | **85871** | **17224** |

**Supplementary Table S6. ATAC-seq quality-control summary.** Per-library read counts, mapping rates, duplication, FRiP scores, and number of peaks called.

| Category | Control 1 | Control 2 | Control 3 | Control 4 | Drought 1 | Drought 2 | Drought 3 | Drought 4 |
|---|---|---|---|---|---|---|---|---|
| **Reads** | 982648218 | 784518374 | 728475838 | 760767412 | 353813162 | 673234566 | 857781406 | 645435518 |
| **Mapped reads** | 965512788 | 763043025 | 703396463 | 741622339 | 337940288 | 639122697 | 685592644 | 441425250 |
| **Mapped and paired reads** | 963789026 | 761277328 | 700347466 | 740225718 | 335332752 | 633660530 | 680053012 | 438280522 |
| **% Mapped reads** | 98,26 | 97,26 | 96,56 | 97,48 | 95,51 | 94,93 | 79,93 | 68,39 |
| **FRIP score** | 0,37 | 0,29 | 0,304 | 0,34 | 0,28 | 0,34 | 0,18 | 0,17 |

**Supplementary Table S7. MethylC-seq quality-control summary.** Per-library read counts, mapping rates, bisulfite conversion rate (estimated from the unmethylated chloroplast control), mean depth per covered cytosine, and genome-wide weighted methylation levels in the CG, CHG, and CHH contexts.

| Read file | % Dups (before trim) | % GC | M Seqs | Uniquely mapped (exactly 1×) | % unique | Multi-mapped (>1×) | Overall alignment (%) |
|---|---|---|---|---|---|---|---|
| control_rep1_R1 | 40,9 | 23 | 421,6 | 139 626 992 | 33,12 | 119 022 194 | 61,35 |
| control_rep1_R2 | 39,9 | 23 | 421,6 | 139 647 036 | 33,12 | 118 699 622 | 61,27 |
| control_rep2_R1 | 41,2 | 23 | 472,4 | 160 028 136 | 33,88 | 136 895 890 | 62,85 |
| control_rep2_R2 | 40,8 | 23 | 472,4 | 160 033 041 | 33,88 | 136 566 502 | 62,79 |
| control_rep3_R1 | 46,0 | 23 | 490,5 | 147 126 891 | 30,00 | 148 832 455 | 60,34 |
| control_rep3_R2 | 45,3 | 23 | 490,5 | 147 326 758 | 30,04 | 148 217 131 | 60,26 |
| drought_rep38_R1 | 42,0 | 23 | 481,3 | 149 730 758 | 31,11 | 141 445 310 | 60,50 |
| drought_rep38_R2 | 41,1 | 23 | 481,3 | 149 765 717 | 31,12 | 140 976 815 | 60,41 |
| drought_rep41_R1 | 40,5 | 23 | 375,7 | 127 117 204 | 33,84 | 105 376 913 | 61,89 |
| drought_rep41_R2 | 39,8 | 23 | 375,7 | 127 131 222 | 33,84 | 105 094 999 | 61,82 |
| drought_rep44_R1 | 44,1 | 23 | 575,6 | 190 537 119 | 33,10 | 154 954 009 | 60,02 |
| drought_rep44_R2 | 43,9 | 23 | 575,6 | 190 508 854 | 33,10 | 154 599 991 | 59,96 |

| Sample | Condition | Total input read pairs | Uniquely mapped read pairs | % uniquely mapped | Non-clonal pairs (post-dedup) | % non-clonal (usable) | Overall alignment rate (%) | Non-conversion rate (%) | Conversion rate (%) |
|---|---|---|---|---|---|---|---|---|---|
| **Control 1 (rep1)** | Well-watered | 421 643 581 | 275 304 363 | 65,3 | 191 667 478 | 45,5 | 61,3 | 0,568 | 99,43 |
| **Control 2 (rep2)** | Well-watered | 472 410 291 | 340 362 692 | 72,0 | 239 597 806 | 50,7 | 62,8 | 0,397 | 99,60 |
| **Control 3 (rep3)** | Well-watered | 490 461 926 | 265 881 207 | 54,2 | 170 427 520 | 34,7 | 60,3 | 0,353 | 99,65 |
| **Drought 1 (rep38)** | Drought | 481 306 489 | 283 563 427 | 58,9 | 192 263 931 | 39,9 | 60,5 | 0,384 | 99,62 |
| **Drought 2 (rep41)** | Drought | 375 683 645 | 269 924 801 | 71,8 | 181 246 374 | 48,2 | 61,9 | 0,431 | 99,57 |
| **Drought 3 (rep44)** | Drought | 575 610 156 | 377 568 587 | 65,6 | 248 622 451 | 43,2 | 60,0 | 0,636 | 99,36 |

**Supplementary Table S8. HOMER motif-enrichment results.** Complete known and de novo motif-enrichment output for the drought-upregulated filtered intergenic dREG peaks, and for the proximal and distal subsets separately, including motif, best match, p-value, q-value, and target/background percentages.

***Known & de novo motif enrichment in upregulated dREGs under drought conditions.***

##### Intergenic known motifs

| Rank | Motif | Name | P-value | log P-pvalue | q-value (Benjamini) | # Target Sequences with Motif | % of Targets Sequences with Motif | # Background Sequences with Motif | % of Background Sequences with Motif | Motif File | SVG |
|---|---|---|---|---|---|---|---|---|---|---|---|
| 1 | 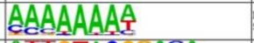 | REM19(REM)/colamp-REM19-DAP-Seq(GSE60143)/Homer | 1e-9 | -2.138e+01 | 0.0000 | 231.0 | 27.47% | 8961.5 | 18.78% | motif file (matrix) | svg |
| 2 | 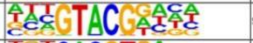 | SPL5(SBP)/colamp-SPL5-DAP-Seq(GSE60143)/Homer | 1e-4 | -1.047e+01 | 0.0071 | 136.0 | 16.17% | 5472.6 | 11.47% | motif file (matrix) | svg |
| 3 | 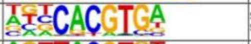 | SPCH(bHLH)/Seedling-SPCH-ChIP-Seq(GSE57497)/Homer | 1e-4 | -1.045e+01 | 0.0071 | 153.0 | 18.19% | 6311.5 | 13.23% | motif file (matrix) | svg |
| 4 | 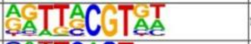 | NAM(NAC)/col-NAM-DAP-Seq(GSE60143)/Homer | 1e-4 | -1.023e+01 | 0.0071 | 181.0 | 21.52% | 7744.4 | 16.23% | motif file (matrix) | svg |
| 5 | 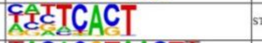 | STZ(C2H2)/colamp-STZ-DAP-Seq(GSE60143)/Homer | 1e-4 | -9.369e+00 | 0.0086 | 447.0 | 53.15% | 22243.7 | 46.62% | motif file (matrix) | svg |
| 6 | 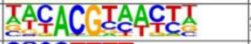 | ANAC047(NAC)/colamp-ANAC047-DAP-Seq(GSE60143)/Homer | 1e-4 | -9.331e+00 | 0.0086 | 117.0 | 13.91% | 4681.6 | 9.81% | motif file (matrix) | svg |
| 7 | 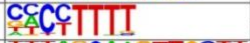 | AT5G02460(C2C2dof)/col-AT5G02460-DAP-Seq(GSE60143)/Homer | 1e-3 | -9.071e+00 | 0.0086 | 293.0 | 34.84% | 13801.8 | 28.93% | motif file (matrix) | svg |
| 8 | 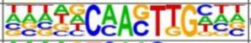 | bHLH80(bHLH)/col-bHLH80-DAP-Seq(GSE60143)/Homer | 1e-3 | -7.864e+00 | 0.0241 | 87.0 | 10.34% | 3404.9 | 7.14% | motif file (matrix) | svg |
| 9 | 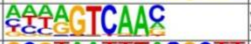 | WRKY40(WRKY)/colamp-WRKY40-DAP-Seq(GSE60143)/Homer | 1e-2 | -6.469e+00 | 0.0865 | 65.0 | 7.73% | 2511.2 | 5.26% | motif file (matrix) | svg |
| 10 | 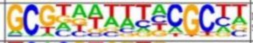 | LBD23(LOBAS2)/colamp-LBD23-DAP-Seq(GSE60143)/Homer | 1e-2 | -6.315e+00 | 0.0908 | 124.0 | 14.74% | 5434.7 | 11.39% | motif file (matrix) | svg |
| 11 | 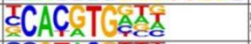 | PIF5ox(bHLH)/Arabidopsis-PIF5ox-ChIP-Seq(GSE35062)/Homer | 1e-2 | -6.172e+00 | 0.0953 | 112.0 | 13.32% | 4850.2 | 10.16% | motif file (matrix) | svg |
| 12 | 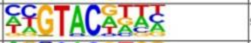 | SPL1(SBP)/colamp-SPL1-DAP-Seq(GSE60143)/Homer | 1e-2 | -5.921e+00 | 0.1122 | 284.0 | 33.77% | 13975.3 | 29.29% | motif file (matrix) | svg |
| 13 | 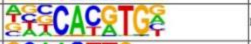 | PIF4(bHLH)/Seedling-PIF4-ChIP-Seq(GSE35315)/Homer | 1e-2 | -5.809e+00 | 0.1158 | 131.0 | 15.58% | 5874.2 | 12.31% | motif file (matrix) | svg |
| 14 | 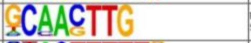 | bHLH130(bHLH)/col-bHLH130-DAP-Seq(GSE60143)/Homer | 1e-2 | -5.714e+00 | 0.1184 | 70.0 | 8.32% | 2836.3 | 5.94% | motif file (matrix) | svg |
| 15 | 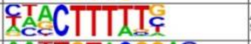 | AT1G47655(C2C2dof)/colamp-AT1G47655-DAP-Seq(GSE60143)/Homer | 1e-2 | -5.647e+00 | 0.1184 | 410.0 | 48.75% | 21027.7 | 44.07% | motif file (matrix) | svg |
| 16 | 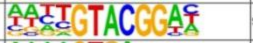 | SPL13(SBP)/col-SPL13-DAP-Seq(GSE60143)/Homer | 1e-2 | -5.562e+00 | 0.1205 | 26.0 | 3.09% | 824.7 | 1.73% | motif file (matrix) | svg |
| 17 | 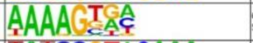 | CDF3(C2C2dof)/colamp-CDF3-DAP-Seq(GSE60143)/Homer | 1e-2 | -5.552e+00 | 0.1205 | 212.0 | 25.21% | 10174.7 | 21.32% | motif file (matrix) | svg |
| 18 | 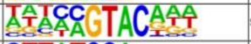 | SPL15(SBP)/colamp-SPL15-DAP-Seq(GSE60143)/Homer | 1e-2 | -5.321e+00 | 0.1363 | 234.0 | 27.82% | 11408.5 | 23.91% | motif file (matrix) | svg |
| 19 | 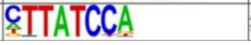 | At5g47390(MYBrelated)/col-At5g47390-DAP-Seq(GSE60143)/Homer | 1e-2 | -5.147e+00 | 0.1537 | 154.0 | 18.31% | 7190.3 | 15.07% | motif file (matrix) | svg |
| 20 | 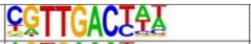 | WRKY24(WRKY)/colamp-WRKY24-DAP-Seq(GSE60143)/Homer | 1e-2 | -5.120e+00 | 0.1537 | 83.0 | 9.87% | 3554.5 | 7.45% | motif file (matrix) | svg |
| 21 | 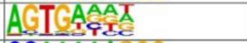 | At5g04390(C2H2)/col200-At5g04390-DAP-Seq(GSE60143)/Homer | 1e-2 | -5.003e+00 | 0.1605 | 431.0 | 51.25% | 22393.3 | 46.93% | motif file (matrix) | svg |
| 22 | 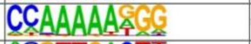 | SEP3(MADS)/Arabidopsis-Flower-Sep3-ChIP-Seq/Homer | 1e-2 | -4.874e+00 | 0.1743 | 159.0 | 18.91% | 7509.3 | 15.74% | motif file (matrix) | svg |
| 23 | 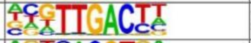 | WRKY55(WRKY)/col-WRKY55-DAP-Seq(GSE60143)/Homer | 1e-2 | -4.767e+00 | 0.1857 | 101.0 | 12.01% | 4518.6 | 9.47% | motif file (matrix) | svg |
| 24 | 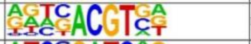 | HY5(bZIP)/colamp-HY5-DAP-Seq(GSE60143)/Homer | 1e-2 | -4.749e+00 | 0.1857 | 130.0 | 15.46% | 6016.4 | 12.61% | motif file (matrix) | svg |
| 25 | 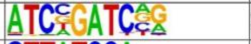 | GATA19(C2C2gata)/colamp-GATA19-DAP-Seq(GSE60143)/Homer | 1e-2 | -4.737e+00 | 0.1857 | 23.0 | 2.73% | 751.2 | 1.57% | motif file (matrix) | svg |
| 26 | 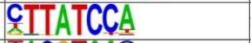 | AT5G61620(MYBrelated)/colamp-AT5G61620-DAP-Seq(GSE60143)/Homer | 1e-2 | -4.711e+00 | 0.1857 | 168.0 | 19.98% | 8020.3 | 16.81% | motif file (matrix) | svg |
| 27 | 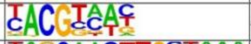 | ATAF1(NAC)/col-ATAF1-DAP-Seq(GSE60143)/Homer | 1e-2 | -4.659e+00 | 0.1857 | 332.0 | 39.48% | 16949.9 | 35.52% | motif file (matrix) | svg |
| 28 | 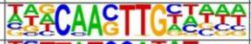 | bHLH122(bHLH)/col100-bHLH122-DAP-Seq(GSE60143)/Homer | 1e-2 | -4.657e+00 | 0.1857 | 77.0 | 9.16% | 3323.1 | 6.96% | motif file (matrix) | svg |
| 29 | 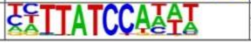 | At1g74840(MYBrelated)/col100-At1g74840-DAP-Seq(GSE60143)/Homer | 1e-2 | -4.646e+00 | 0.1857 | 113.0 | 13.44% | 5155.4 | 10.80% | motif file (matrix) | svg |

#### Intergenic *de novo* motifs

| Rank | Motif | P-value | log P-pvalue | % of Targets | % of Background | STD(Bg STD) | Best Match/Details | Motif File |
|---|---|---|---|---|---|---|---|---|
| 1 | 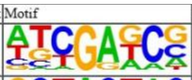 | 1e-17 | -4.131e+01 | 18.19% | 8.59% | 57.1bp (67.4bp) | GATA20(C2C2gata)/colamp-GATA20-DAP-Seq(GSE60143)/Homer(0.744) More Information \| Similar Motifs Found | motif file (matrix) |
| 2 | 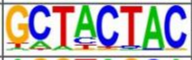 | 1e-14 | -3.326e+01 | 17.12% | 8.64% | 53.4bp (63.3bp) | At3g60580(C2H2)/col-At3g60580-DAP-Seq(GSE60143)/Homer(0.614) More Information \| Similar Motifs Found | motif file (matrix) |
| 3 | 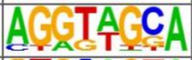 | 1e-12 | -2.916e+01 | 12.72% | 5.95% | 55.0bp (60.3bp) | P(MYB)/Zea mays/AthaMap(0.611) More Information \| Similar Motifs Found | motif file (matrix) |
| 4 | 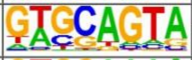 | 1e-12 | -2.900e+01 | 39.83% | 28.19% | 54.8bp (62.4bp) | At3g60580(C2H2)/col-At3g60580-DAP-Seq(GSE60143)/Homer(0.695) More Information \| Similar Motifs Found | motif file (matrix) |
| 5 | 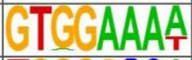 | 1e-12 | -2.807e+01 | 9.63% | 3.98% | 53.0bp (61.0bp) | At5g47390(MYBrelated)/col-At5g47390-DAP-Seq(GSE60143)/Homer(0.704) More Information \| Similar Motifs Found | motif file (matrix) |
| 6 | 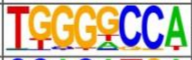 | 1e-12 | -2.768e+01 | 11.41% | 5.19% | 57.5bp (58.5bp) | TCP2/MA1064.1/Jaspar(0.822) More Information \| Similar Motifs Found | motif file (matrix) |
| 7 * | 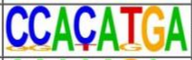 | 1e-11 | -2.732e+01 | 7.97% | 3.00% | 53.8bp (60.9bp) | PIF4(bHLH)/Seedling-PIF4-ChIP-Seq(GSE35315)/Homer(0.758) More Information \| Similar Motifs Found | motif file (matrix) |
| 8 * | 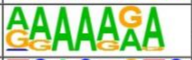 | 1e-11 | -2.713e+01 | 38.17% | 27.07% | 53.8bp (69.4bp) | REM19(REM)/colamp-REM19-DAP-Seq(GSE60143)/Homer(0.844) More Information \| Similar Motifs Found | motif file (matrix) |
| 9 * | 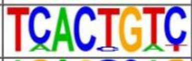 | 1e-10 | -2.469e+01 | 7.49% | 2.90% | 55.8bp (60.5bp) | At5g04390(C2H2)/col200-At5g04390-DAP-Seq(GSE60143)/Homer(0.772) More Information \| Similar Motifs Found | motif file (matrix) |
| 10 * | 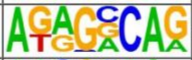 | 1e-9 | -2.277e+01 | 16.77% | 9.69% | 49.9bp (58.5bp) | NLP7(RWPRK)/col-NLP7-DAP-Seq(GSE60143)/Homer(0.740) More Information \| Similar Motifs Found | motif file (matrix) |
| 11 * | 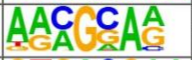 | 1e-9 | -2.209e+01 | 34.01% | 24.40% | 57.6bp (62.1bp) | Dof2/MA0020.1/Jaspar(0.712) More Information \| Similar Motifs Found | motif file (matrix) |
| 12 * | 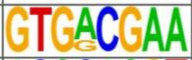 | 1e-9 | -2.186e+01 | 4.04% | 1.12% | 54.4bp (70.7bp) | TGA2(bZIP)/colamp-TGA2-DAP-Seq(GSE60143)/Homer(0.731) More Information \| Similar Motifs Found | motif file (matrix) |
| 13 * | 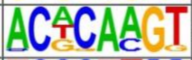 | 1e-9 | -2.093e+01 | 7.49% | 3.19% | 51.7bp (66.4bp) | BHLH112/MA0961.1/Jaspar(0.695) More Information \| Similar Motifs Found | motif file (matrix) |
| 14 * | 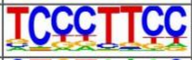 | 1e-8 | -1.967e+01 | 35.32% | 26.15% | 58.9bp (58.7bp) | Knotted(Homeobox)/Corn-KN1-ChIP-Seq(GSE39161)/Homer(0.702) More Information \| Similar Motifs Found | motif file (matrix) |
| 15 * | 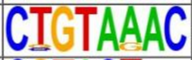 | 1e-7 | -1.633e+01 | 3.45% | 1.08% | 46.0bp (58.8bp) | GTL1(Trihelix)/colamp-GTL1-DAP-Seq(GSE60143)/Homer(0.661) More Information \| Similar Motifs Found | motif file (matrix) |
| 16 * | 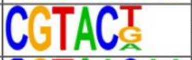 | 1e-7 | -1.618e+01 | 14.03% | 8.55% | 58.0bp (73.3bp) | SPL5(SBP)/colamp-SPL5-DAP-Seq(GSE60143)/Homer(0.958) More Information \| Similar Motifs Found | motif file (matrix) |
| 17 * | 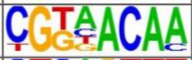 | 1e-7 | -1.614e+01 | 7.37% | 3.55% | 57.3bp (62.2bp) | MF0011.1_HMG_class/Jaspar(0.751) More Information \| Similar Motifs Found | motif file (matrix) |
| 18 * | 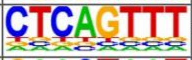 | 1e-6 | -1.557e+01 | 2.50% | 0.63% | 35.9bp (57.5bp) | JGL(C2H2)/col-JGL-DAP-Seq(GSE60143)/Homer(0.699) More Information \| Similar Motifs Found | motif file (matrix) |
| 19 * | 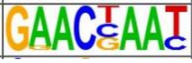 | 1e-6 | -1.489e+01 | 7.13% | 3.52% | 55.1bp (65.0bp) | MYB107(MYB)/col-MYB107-DAP-Seq(GSE60143)/Homer(0.651) More Information \| Similar Motifs Found | motif file (matrix) |
| 20 * | 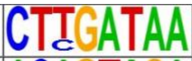 | 1e-6 | -1.483e+01 | 2.97% | 0.90% | 45.2bp (56.9bp) | At5g47390(MYBrelated)/col-At5g47390-DAP-Seq(GSE60143)/Homer(0.770) More Information \| Similar Motifs Found | motif file (matrix) |
| 21 * | 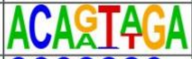 | 1e-4 | -1.046e+01 | 7.85% | 4.62% | 55.4bp (58.0bp) | MF0011.1_HMG_class/Jaspar(0.645) More Information \| Similar Motifs Found | motif file (matrix) |
| 22 * | 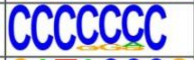 | 1e-4 | -1.009e+01 | 11.41% | 7.54% | 49.3bp (57.7bp) | POL003.1_GC-box/Jaspar(0.706) More Information \| Similar Motifs Found | motif file (matrix) |
| 23 * | 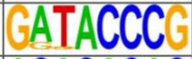 | 1e-4 | -9.243e+00 | 1.90% | 0.62% | 47.3bp (61.7bp) | CMTA2/MA0969.1/Jaspar(0.618) More Information \| Similar Motifs Found | motif file (matrix) |
| 24 * | 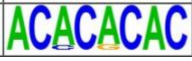 | 1e-1 | -4.327e+00 | 4.04% | 2.68% | 71.9bp (63.8bp) | NAC080/MA0939.1/Jaspar(0.605) More Information \| Similar Motifs Found | motif file (matrix) |

#### Distal known motifs

| Rank | Motif | Name | P-value | log P-pvalue | q-value (Benjamini) | # Target Sequences with Motif | % of Targets Sequences with Motif | # Background Sequences with Motif | % of Background Sequences with Motif | Motif File | SVG |
|---|---|---|---|---|---|---|---|---|---|---|---|
| 1 | 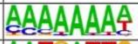 | REM19(REM)/colamp-REM19-DAP-Seq(GSE60143)/Homer | 1e-5 | -1.192e+01 | 0.0033 | 103.0 | 32.09% | 10308.6 | 21.48% | motif file (matrix) | svg |
| 2 | 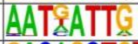 | ATHB5(HB)/colamp-ATHB5-DAP-Seq(GSE60143)/Homer | 1e-3 | -8.357e+00 | 0.0589 | 82.0 | 25.55% | 8452.6 | 17.61% | motif file (matrix) | svg |
| 3 | 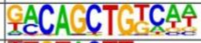 | bZIP69(bZIP)/col-bZIP69-DAP-Seq(GSE60143)/Homer | 1e-3 | -7.018e+00 | 0.1499 | 6.0 | 1.87% | 163.8 | 0.34% | motif file (matrix) | svg |
| 4 | 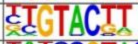 | SPL9(SBP)/colamp-SPL9-DAP-Seq(GSE60143)/Homer | 1e-2 | -6.759e+00 | 0.1499 | 117.0 | 36.45% | 13659.4 | 28.46% | motif file (matrix) | svg |
| 5 | 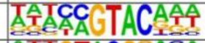 | SPL15(SBP)/colamp-SPL15-DAP-Seq(GSE60143)/Homer | 1e-2 | -6.266e+00 | 0.1907 | 96.0 | 29.91% | 10937.0 | 22.79% | motif file (matrix) | svg |
| 6 | 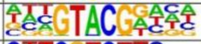 | SPL5(SBP)/colamp-SPL5-DAP-Seq(GSE60143)/Homer | 1e-2 | -6.146e+00 | 0.1907 | 52.0 | 16.20% | 5194.4 | 10.82% | motif file (matrix) | svg |
| 7 | 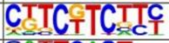 | Unknown4/Arabidopsis-Promoters/Homer | 1e-2 | -6.099e+00 | 0.1907 | 65.0 | 20.25% | 6863.4 | 14.30% | motif file (matrix) | svg |
| 8 | 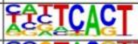 | STZ(C2H2)/colamp-STZ-DAP-Seq(GSE60143)/Homer | 1e-2 | -5.645e+00 | 0.2219 | 172.0 | 53.58% | 22042.5 | 45.92% | motif file (matrix) | svg |
| 9 | 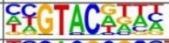 | SPL1(SBP)/colamp-SPL1-DAP-Seq(GSE60143)/Homer | 1e-2 | -5.635e+00 | 0.2219 | 111.0 | 34.58% | 13239.1 | 27.58% | motif file (matrix) | svg |
| 10 | 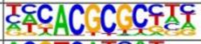 | FHY3(FAR1)/Arabidopsis-FHY3-ChIP-Seq(GSE30711)/Homer | 1e-2 | -4.898e+00 | 0.3745 | 25.0 | 7.79% | 2198.5 | 4.58% | motif file (matrix) | svg |
| 11 | 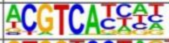 | TGA2(bZIP)/colamp-TGA2-DAP-Seq(GSE60143)/Homer | 1e-2 | -4.663e+00 | 0.4306 | 43.0 | 13.40% | 4446.1 | 9.26% | motif file (matrix) | svg |
| 12 | 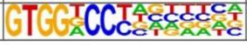 | TCP16(TCP)/colamp-TCP16-DAP-Seq(GSE60143)/Homer | 1e-2 | -4.656e+00 | 0.4306 | 18.0 | 5.61% | 1447.5 | 3.02% | motif file (matrix) | svg |

#### Distal *de novo* motifs

| Rank | Motif | P-value | log P-pvalue | % of Targets | % of Background | STD(Bg STD) | Best Match/Details | Motif File |
|---|---|---|---|---|---|---|---|---|
| 1 | CTGGACCG | 1e-14 | -3.450e+01 | 6.85% | 0.66% | 54.3bp (63.8bp) | TCP16/MA0587.1/Jaspar(0.681)<br>More Information \| Similar Motifs Found | motif file (matrix) |
| 2 * | TGCCTATC | 1e-11 | -2.749e+01 | 13.71% | 3.96% | 52.2bp (59.6bp) | MYB116(MYB)/colamp-MYB116-DAP-Seq(GSE60143)/Homer(0.723)<br>More Information \| Similar Motifs Found | motif file (matrix) |
| 3 * | GGAAAAAA | 1e-9 | -2.276e+01 | 12.77% | 4.06% | 56.5bp (68.9bp) | REM19(REM)/colamp-REM19-DAP-Seq(GSE60143)/Homer(0.823)<br>More Information \| Similar Motifs Found | motif file (matrix) |
| 4 * | GGTTGCGT | 1e-9 | -2.154e+01 | 11.21% | 3.37% | 52.6bp (62.8bp) | NAC083/MA1043.1/Jaspar(0.823)<br>More Information \| Similar Motifs Found | motif file (matrix) |
| 5 * | TGGCGAC | 1e-9 | -2.094e+01 | 13.08% | 4.50% | 47.4bp (64.7bp) | ERF105(AP2EREBP)/colamp-ERF105-DAP-Seq(GSE60143)/Homer(0.725) <br>More Information \| Similar Motifs Found | motif file (matrix) |
| 6 * | TATGTAG | 1e-8 | -1.938e+01 | 22.74% | 11.25% | 54.7bp (59.5bp) | DDF1(AP2EREBP)/col-DDF1-DAP-Seq(GSE60143)/Homer(0.708)<br>More Information \| Similar Motifs Found | motif file (matrix) |
| 7 * | ATACCGCA | 1e-7 | -1.826e+01 | 6.23% | 1.29% | 57.9bp (61.8bp) | AT1G76870(Trihelix)/col-AT1G76870-DAP-Seq(GSE60143)/Homer(0.586)<br>More Information \| Similar Motifs Found | motif file (matrix) |
| 8 * | GCTCTATC | 1e-7 | -1.810e+01 | 14.02% | 5.55% | 44.6bp (59.9bp) | POL009.1_DCE_S_II/Jaspar(0.585)<br>More Information \| Similar Motifs Found | motif file (matrix) |
| 9 * | GGGTCAAC | 1e-7 | -1.730e+01 | 8.72% | 2.58% | 56.2bp (57.1bp) | WRKY63/MA1092.1/Jaspar(0.889)<br>More Information \| Similar Motifs Found | motif file (matrix) |
| 10 * | TCGTCGT | 1e-6 | -1.588e+01 | 17.76% | 8.57% | 52.4bp (70.1bp) | MGP(C2H2)/colamp-MGP-DAP-Seq(GSE60143)/Homer(0.718)<br>More Information \| Similar Motifs Found | motif file (matrix) |
| 11 * | ATCGAGA | 1e-6 | -1.409e+01 | 7.48% | 2.33% | 54.7bp (63.1bp) | GATA20(C2C2gata)/colamp-GATA20-DAP-Seq(GSE60143)/Homer(0.663)<br>More Information \| Similar Motifs Found | motif file (matrix) |
| 12 * | ACGAAAAG | 1e-5 | -1.312e+01 | 8.72% | 3.18% | 46.0bp (60.7bp) | CDF3(C2C2dof)/colamp-CDF3-DAP-Seq(GSE60143)/Homer(0.721)<br>More Information \| Similar Motifs Found | motif file (matrix) |
| 13 * | ACTAATTA | 1e-5 | -1.307e+01 | 5.92% | 1.64% | 50.7bp (58.3bp) | ATHB23(ZFHD)/col-ATHB23-DAP-Seq(GSE60143)/Homer(0.872)<br>More Information \| Similar Motifs Found | motif file (matrix) |
| 14 * | GGGCTGC | 1e-5 | -1.284e+01 | 5.61% | 1.51% | 46.9bp (60.9bp) | POL010.1_DCE_S_III/Jaspar(0.721)<br>More Information \| Similar Motifs Found | motif file (matrix) |
| 15 * | GCATGCT | 1e-5 | -1.270e+01 | 11.84% | 5.25% | 50.8bp (65.4bp) | FUS3/MA0565.1/Jaspar(0.923)<br>More Information \| Similar Motifs Found | motif file (matrix) |
| 16 * | AGTGTAC | 1e-5 | -1.232e+01 | 24.92% | 15.25% | 55.5bp (60.7bp) | SPL4/MA1058.1/Jaspar(0.719)<br>More Information \| Similar Motifs Found | motif file (matrix) |
| 17 * | ATGGAAGG | 1e-4 | -1.136e+01 | 4.67% | 1.21% | 52.9bp (58.4bp) | POL008.1_DCE_S_I/Jaspar(0.698)<br>More Information \| Similar Motifs Found | motif file (matrix) |
| 18 * | AACCGATA | 1e-4 | -1.013e+01 | 4.05% | 1.04% | 64.7bp (60.7bp) | MYB70(MYB)/col-MYB70-DAP-Seq(GSE60143)/Homer(0.836)<br>More Information \| Similar Motifs Found | motif file (matrix) |
| 19 * | AAAGACTT | 1e-4 | -9.664e+00 | 5.61% | 1.92% | 56.5bp (56.6bp) | DOF5.7/MA0984.1/Jaspar(0.626)<br>More Information \| Similar Motifs Found | motif file (matrix) |
| 20 * | ATACCCGC | 1e-3 | -8.498e+00 | 1.56% | 0.16% | 28.2bp (49.0bp) | DEL2(E2FDP)/col-DEL2-DAP-Seq(GSE60143)/Homer(0.643)<br>More Information \| Similar Motifs Found | motif file (matrix) |
| 21 * | TTCTCTCA | 1e-2 | -6.838e+00 | 4.67% | 1.84% | 44.3bp (59.7bp) | BPC1(BBRBPC)/colamp-BPC1-DAP-Seq(GSE60143)/Homer(0.752)<br>More Information \| Similar Motifs Found | motif file (matrix) |
| 22 * | CCACGCG | 1e-2 | -5.905e+00 | 4.67% | 2.03% | 48.3bp (68.8bp) | FHY3(FAR1)/Arabidopsis-FHY3-ChIP-Seq(GSE30711)/Homer(0.856)<br>More Information \| Similar Motifs Found | motif file (matrix) |
| 23 * | TTTAAAAA | 1e-1 | -4.570e+00 | 10.59% | 6.97% | 56.3bp (69.1bp) | At2g41835(C2H2)/col-At2g41835-DAP-Seq(GSE60143)/Homer(0.807)<br>More Information \| Similar Motifs Found | motif file (matrix) |
| 24 * | ATAATAA | 1e-1 | -3.934e+00 | 14.02% | 10.25% | 51.3bp (58.6bp) | KANADI1(Myb)/Seedling-KAN1-ChIP-Seq(GSE48081)/Homer(0.766)<br>More Information \| Similar Motifs Found | motif file (matrix) |

#### Proximal intergenic known motifs

| Rank | Motif | Name | P-value | log P-pvalue | q-value (Benjamini) | # Target Sequences with Motif | % of Targets Sequences with Motif | # Background Sequences with Motif | % of Background Sequences with Motif | Motif File | SVG |
|---|---|---|---|---|---|---|---|---|---|---|---|
| 1 | 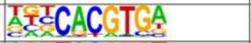 | SPCH(bHLH)/Seedling-SPCH-ChIP-Seq(GSE57497)/Homer | 1e-4 | -1.115e+01 | 0.0072 | 106.0 | 20.38% | 6546.3 | 13.64% | motif file (matrix) | svg |
| 2 | 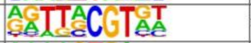 | NAM(NAC)/col-NAM-DAP-Seq(GSE60143)/Homer | 1e-4 | -1.086e+01 | 0.0072 | 123.0 | 23.65% | 7937.9 | 16.54% | motif file (matrix) | svg |
| 3 | 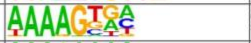 | CDF3(C2C2dof)/colamp-CDF3-DAP-Seq(GSE60143)/Homer | 1e-4 | -1.057e+01 | 0.0072 | 146.0 | 28.08% | 9859.6 | 20.54% | motif file (matrix) | svg |
| 4 | 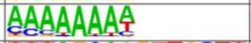 | REM19(REM)/colamp-REM19-DAP-Seq(GSE60143)/Homer | 1e-4 | -1.044e+01 | 0.0072 | 128.0 | 24.62% | 8413.0 | 17.53% | motif file (matrix) | svg |
| 5 | 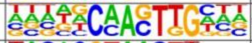 | bHLH80(bHLH)/col-bHLH80-DAP-Seq(GSE60143)/Homer | 1e-3 | -8.583e+00 | 0.0188 | 61.0 | 11.73% | 3496.0 | 7.28% | motif file (matrix) | svg |
| 6 | 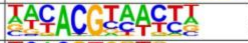 | ANAC047(NAC)/colamp-ANAC047-DAP-Seq(GSE60143)/Homer | 1e-3 | -8.334e+00 | 0.0201 | 76.0 | 14.62% | 4663.3 | 9.72% | motif file (matrix) | svg |
| 7 | 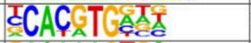 | PIF5ox(bHLH)/Arabidopsis-PIF5ox-ChIP-Seq(GSE35062)/Homer | 1e-3 | -8.101e+00 | 0.0217 | 81.0 | 15.58% | 5087.1 | 10.60% | motif file (matrix) | svg |
| 8 | 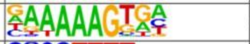 | COG1(C2C2dof)/col-COG1-DAP-Seq(GSE60143)/Homer | 1e-3 | -7.881e+00 | 0.0237 | 129.0 | 24.81% | 9001.4 | 18.75% | motif file (matrix) | svg |
| 9 | 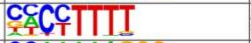 | AT5G02460(C2C2dof)/col-AT5G02460-DAP-Seq(GSE60143)/Homer | 1e-3 | -7.765e+00 | 0.0237 | 181.0 | 34.81% | 13445.8 | 28.01% | motif file (matrix) | svg |
| 10 | 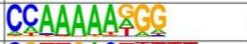 | SEP3(MADS)/Arabidoposis-Flower-Sep3-ChIP-Seq/Homer | 1e-3 | -7.126e+00 | 0.0403 | 106.0 | 20.38% | 7268.7 | 15.14% | motif file (matrix) | svg |
| 11 | 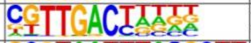 | WRKY26(WRKY)/colamp-WRKY26-DAP-Seq(GSE60143)/Homer | 1e-3 | -6.987e+00 | 0.0422 | 39.0 | 7.50% | 2100.9 | 4.38% | motif file (matrix) | svg |
| 12 | 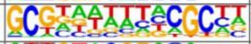 | LBD23(LOBAS2)/colamp-LBD23-DAP-Seq(GSE60143)/Homer | 1e-2 | -6.900e+00 | 0.0422 | 85.0 | 16.35% | 5616.3 | 11.70% | motif file (matrix) | svg |
| 13 | 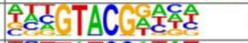 | SPL5(SBP)/colamp-SPL5-DAP-Seq(GSE60143)/Homer | 1e-2 | -6.540e+00 | 0.0558 | 84.0 | 16.15% | 5606.8 | 11.68% | motif file (matrix) | svg |
| 14 | 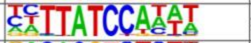 | At1g74840(MYBrelated)/col100-At1g74840-DAP-Seq(GSE60143)/Homer | 1e-2 | -6.495e+00 | 0.0558 | 77.0 | 14.81% | 5056.0 | 10.53% | motif file (matrix) | svg |
| 15 | 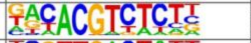 | bHLH157(bHLH)/col-bHLH157-DAP-Seq(GSE60143)/Homer | 1e-2 | -6.413e+00 | 0.0558 | 24.0 | 4.62% | 1129.2 | 2.35% | motif file (matrix) | svg |
| 16 | 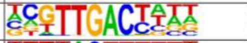 | WRKY6(WRKY)/colamp-WRKY6-DAP-Seq(GSE60143)/Homer | 1e-2 | -6.152e+00 | 0.0668 | 58.0 | 11.15% | 3629.1 | 7.56% | motif file (matrix) | svg |
| 17 | 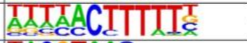 | OBP4(C2C2dof)/col-OBP4-DAP-Seq(GSE60143)/Homer | 1e-2 | -6.150e+00 | 0.0668 | 133.0 | 25.58% | 9749.5 | 20.31% | motif file (matrix) | svg |
| 18 | 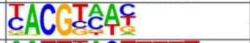 | ATAF1(NAC)/col-ATAF1-DAP-Seq(GSE60143)/Homer | 1e-2 | -6.004e+00 | 0.0688 | 220.0 | 42.31% | 17391.7 | 36.23% | motif file (matrix) | svg |
| 19 | 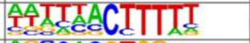 | DAG2(C2C2dof)/col-DAG2-DAP-Seq(GSE60143)/Homer | 1e-2 | -6.001e+00 | 0.0688 | 148.0 | 28.46% | 11070.9 | 23.06% | motif file (matrix) | svg |
| 20 | 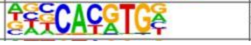 | PIF4(bHLH)/Seedling-PIF4-ChIP-Seq(GSE35315)/Homer | 1e-2 | -5.838e+00 | 0.0731 | 89.0 | 17.12% | 6157.8 | 12.83% | motif file (matrix) | svg |
| 21 | 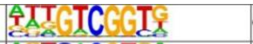 | CEJ1(AP2EREBP)/col-CEJ1-DAP-Seq(GSE60143)/Homer | 1e-2 | -5.679e+00 | 0.0817 | 94.0 | 18.08% | 6603.2 | 13.76% | motif file (matrix) | svg |
| 22 | 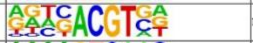 | HY5(bZIP)/colamp-HY5-DAP-Seq(GSE60143)/Homer | 1e-2 | -5.453e+00 | 0.0977 | 89.0 | 17.12% | 6244.7 | 13.01% | motif file (matrix) | svg |
| 23 | 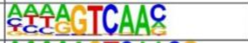 | WRKY40(WRKY)/colamp-WRKY40-DAP-Seq(GSE60143)/Homer | 1e-2 | -5.306e+00 | 0.1082 | 41.0 | 7.88% | 2467.3 | 5.14% | motif file (matrix) | svg |
| 24 | 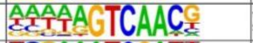 | WRKY65(WRKY)/colamp-WRKY65-DAP-Seq(GSE60143)/Homer | 1e-2 | -5.287e+00 | 0.1082 | 37.0 | 7.12% | 2171.1 | 4.52% | motif file (matrix) | svg |
| 25 | 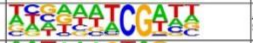 | AT2G15740(C2H2)/col-AT2G15740-DAP-Seq(GSE60143)/Homer | 1e-2 | -5.202e+00 | 0.1105 | 157.0 | 30.19% | 12084.1 | 25.18% | motif file (matrix) | svg |
| 26 | 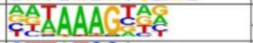 | Adof1(C2C2dof)/col-Adof1-DAP-Seq(GSE60143)/Homer | 1e-2 | -5.192e+00 | 0.1105 | 203.0 | 39.04% | 16142.7 | 33.63% | motif file (matrix) | svg |
| 27 | 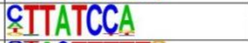 | At5g47390(MYBrelated)/col-At5g47390-DAP-Seq(GSE60143)/Homer | 1e-2 | -5.185e+00 | 0.1105 | 99.0 | 19.04% | 7137.1 | 14.87% | motif file (matrix) | svg |
| 28 | 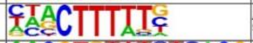 | AT1G47655(C2C2dof)/colamp-AT1G47655-DAP-Seq(GSE60143)/Homer | 1e-2 | -5.158e+00 | 0.1105 | 250.0 | 48.08% | 20391.9 | 42.48% | motif file (matrix) | svg |
| 29 | 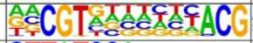 | ANAC094(NAC)/col-ANAC094-DAP-Seq(GSE60143)/Homer | 1e-2 | -5.123e+00 | 0.1105 | 50.0 | 9.62% | 3185.3 | 6.64% | motif file (matrix) | svg |
| 30 | 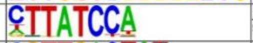 | AT5G61620(MYBrelated)/colamp-AT5G61620-DAP-Seq(GSE60143)/Homer | 1e-2 | -4.943e+00 | 0.1193 | 108.0 | 20.77% | 7956.2 | 16.58% | motif file (matrix) | svg |
| 31 | 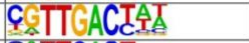 | WRKY24(WRKY)/colamp-WRKY24-DAP-Seq(GSE60143)/Homer | 1e-2 | -4.865e+00 | 0.1249 | 53.0 | 10.19% | 3467.0 | 7.22% | motif file (matrix) | svg |
| 32 | 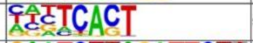 | STZ(C2H2)/colamp-STZ-DAP-Seq(GSE60143)/Homer | 1e-2 | -4.709e+00 | 0.1414 | 275.0 | 52.88% | 22850.3 | 47.61% | motif file (matrix) | svg |
| 33 | 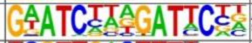 | At1g13300(G2like)/col-At1g13300-DAP-Seq(GSE60143)/Homer | 1e-2 | -4.694e+00 | 0.1414 | 3.0 | 0.58% | 39.5 | 0.08% | motif file (matrix) | svg |
| 34 | 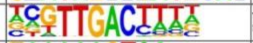 | WRKY14(WRKY)/colamp-WRKY14-DAP-Seq(GSE60143)/Homer | 1e-2 | -4.669e+00 | 0.1414 | 35.0 | 6.73% | 2111.6 | 4.40% | motif file (matrix) | svg |
| 35 | 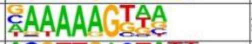 | AT2G28810(C2C2dof)/colamp-AT2G28810-DAP-Seq(GSE60143)/Homer | 1e-2 | -4.654e+00 | 0.1414 | 169.0 | 32.50% | 13315.7 | 27.74% | motif file (matrix) | svg |
| 36 | 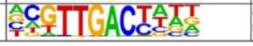 | WRKY31(WRKY)/colamp-WRKY31-DAP-Seq(GSE60143)/Homer | 1e-2 | -4.641e+00 | 0.1414 | 45.0 | 8.65% | 2881.5 | 6.00% | motif file (matrix) | svg |

#### Proximal intergenic *de novo* motifs

| Rank | Motif | P-value | log P-pvalue | % of Targets | % of Background | STD(Bg STD) | Best Match/Details | Motif File |
|---|---|---|---|---|---|---|---|---|
| 1 | 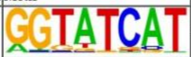 | 1e-14 | -3.378e+01 | 15.96% | 6.12% | 53.9bp (59.1bp) | CRC(C2C2YABBY)/col-CRC-DAP-Seq(GSE60143)/Homer(0.750) More Information \| Similar Motifs Found | motif file (matrix) |
| 2 | 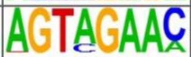 | 1e-13 | -3.150e+01 | 12.50% | 4.27% | 55.4bp (62.2bp) | At3g60580(C2H2)/col-At3g60580-DAP-Seq(GSE60143)/Homer(0.809) More Information \| Similar Motifs Found | motif file (matrix) |
| 3 | 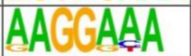 | 1e-12 | -2.826e+01 | 28.65% | 16.12% | 54.3bp (57.5bp) | AT5G60130(ABI3VP1)/col-AT5G60130-DAP-Seq(GSE60143)/Homer(0.797) More Information \| Similar Motifs Found | motif file (matrix) |
| 4 * | 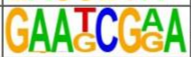 | 1e-11 | -2.694e+01 | 8.65% | 2.53% | 49.7bp (62.8bp) | POL008.1_DCE_S_I/Jaspar(0.652) More Information \| Similar Motifs Found | motif file (matrix) |
| 5 * | 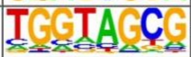 | 1e-11 | -2.684e+01 | 40.38% | 26.32% | 57.5bp (62.1bp) | CRF4(AP2EREBP)/colamp-CRF4-DAP-Seq(GSE60143)/Homer(0.622) More Information \| Similar Motifs Found | motif file (matrix) |
| 6 * | 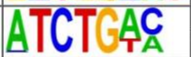 | 1e-11 | -2.627e+01 | 25.58% | 14.10% | 53.9bp (62.1bp) | AtGRF6(GRF)/col-AtGRF6-DAP-Seq(GSE60143)/Homer(0.775) More Information \| Similar Motifs Found | motif file (matrix) |
| 7 * | 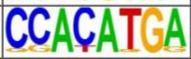 | 1e-11 | -2.619e+01 | 10.58% | 3.66% | 49.4bp (60.9bp) | PIF4(bHLH)/Seedling-PIF4-ChIP-Seq(GSE35315)/Homer(0.781) More Information \| Similar Motifs Found | motif file (matrix) |
| 8 * | 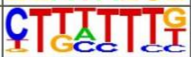 | 1e-11 | -2.543e+01 | 44.81% | 30.68% | 53.4bp (67.1bp) | BBX31(Orphan)/col-BBX31-DAP-Seq(GSE60143)/Homer(0.774) More Information \| Similar Motifs Found | motif file (matrix) |
| 9 * | 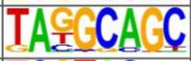 | 1e-10 | -2.506e+01 | 8.65% | 2.68% | 61.5bp (63.8bp) | MYB116(MYB)/colamp-MYB116-DAP-Seq(GSE60143)/Homer(0.676) More Information \| Similar Motifs Found | motif file (matrix) |
| 10 * | 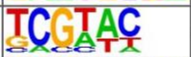 | 1e-10 | -2.456e+01 | 36.15% | 23.22% | 58.4bp (69.1bp) | SPL5(SBP)/colamp-SPL5-DAP-Seq(GSE60143)/Homer(0.838) More Information \| Similar Motifs Found | motif file (matrix) |
| 11 * | 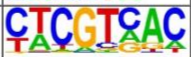 | 1e-9 | -2.212e+01 | 16.15% | 7.82% | 54.1bp (66.5bp) | TGA2(bZIP)/colamp-TGA2-DAP-Seq(GSE60143)/Homer(0.790) More Information \| Similar Motifs Found | motif file (matrix) |
| 12 * | 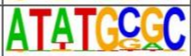 | 1e-9 | -2.149e+01 | 6.54% | 1.85% | 46.6bp (62.1bp) | PCF/Arabidopsis-Promoters/Homer(0.676) More Information \| Similar Motifs Found | motif file (matrix) |
| 13 * | 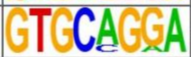 | 1e-8 | -1.896e+01 | 6.15% | 1.84% | 46.8bp (62.8bp) | POL011.1_XCPE1/Jaspar(0.636) More Information \| Similar Motifs Found | motif file (matrix) |
| 14 * | 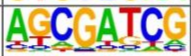 | 1e-8 | -1.883e+01 | 16.92% | 8.96% | 59.2bp (69.6bp) | GATA20(C2C2gata)/colamp-GATA20-DAP-Seq(GSE60143)/Homer(0.708) More Information \| Similar Motifs Found | motif file (matrix) |
| 15 * | 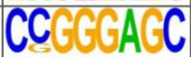 | 1e-8 | -1.856e+01 | 3.85% | 0.77% | 48.3bp (59.9bp) | POL013.1_MED-1/Jaspar(0.805) More Information \| Similar Motifs Found | motif file (matrix) |
| 16 * | 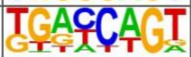 | 1e-7 | -1.755e+01 | 26.54% | 16.92% | 55.2bp (61.5bp) | MYB81(MYB)/col-MYB81-DAP-Seq(GSE60143)/Homer(0.661) More Information \| Similar Motifs Found | motif file (matrix) |
| 17 * | 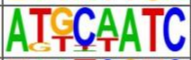 | 1e-7 | -1.643e+01 | 13.27% | 6.73% | 56.3bp (60.0bp) | WUS1(Homeobox)/colamp-WUS1-DAP-Seq(GSE60143)/Homer(0.691) More Information \| Similar Motifs Found | motif file (matrix) |
| 18 * | 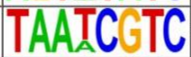 | 1e-7 | -1.639e+01 | 2.69% | 0.42% | 33.1bp (60.1bp) | AtMYB77(MYB)/Arabidopsis thaliana/AthaMap(0.720) More Information \| Similar Motifs Found | motif file (matrix) |
| 19 * | 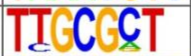 | 1e-7 | -1.625e+01 | 16.15% | 8.90% | 52.6bp (65.4bp) | E2FA(E2FDP)/colamp-E2FA-DAP-Seq(GSE60143)/Homer(0.661) More Information \| Similar Motifs Found | motif file (matrix) |
| 20 * | 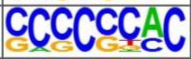 | 1e-6 | -1.574e+01 | 6.35% | 2.24% | 40.9bp (56.6bp) | ALFIN1(HD-PHD)/Medicago sativa/AthaMap(0.730) More Information \| Similar Motifs Found | motif file (matrix) |
| 21 * | 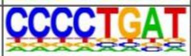 | 1e-6 | -1.556e+01 | 2.12% | 0.26% | 44.6bp (53.2bp) | RAV1(2)(AP2/EREBP)/Arabidopsis thaliana/AthaMap(0.720) More Information \| Similar Motifs Found | motif file (matrix) |
